# Spatial positioning and matrix programs of cancer-associated fibroblasts promote T cell exclusion in human lung tumors

**DOI:** 10.1101/2022.01.20.476763

**Authors:** John A. Grout, Philémon Sirven, Andrew M. Leader, Shrisha Maskey, Eglantine Hector, Isabelle Puisieux, Fiona Steffan, Evan Cheng, Navpreet Tung, Mathieu Maurin, Romain Vaineau, Léa Karpf, Martin Plaud, Maria Casanova-Acebes, Alexandra Tabachnikova, Shilpa Keerthivasan, Alona Lansky, Jessica LeBérichel, Laura Walker, Adeeb H. Rahman, Sacha Gnjatic, Julien Adam, Jerome C. Martin, Andrea Wolf, Raja Flores, Mary Beth Beasley, Rachana Pradhan, Sören Müller, Thomas U. Marron, Shannon J. Turley, Miriam Merad, Ephraim Kenigsberg, Hélène Salmon

## Abstract

It is currently accepted that activated cancer-associated fibroblasts (CAF) participate in T cell exclusion from tumor nests, but it remains unclear how they promote barrier phenotypes, and whether specific subsets are involved. Here, using single-cell RNA sequencing coupled with multiplex imaging on a large cohort of lung tumors, we identify four main CAF populations, of which only two are associated with T cell exclusion: (i) MYH11^+^αSMA^+^ CAF, which are present in early-stage tumors and form a single-cell layer lining cancer aggregates, and (ii) FAP^+^αSMA^+^ CAF, which appear in more advanced tumors and organize in patches within the stroma or in multiple layers around tumor nests. Both CAF populations show a contractility phenotype together with dense and aligned matrix fiber deposition compared to the T cell-permissive CAF. Yet they express distinct matrix genes, including COL4A1/COL9A1 (MYH11^+^αSMA^+^ CAF) and COL11A1/COL12A1 (FAP^+^αSMA^+^ CAF). Hereby, we uncovered unique molecular programs of CAF driving T cell marginalization, whose targeting should increase immunotherapy efficacy in patients bearing T cell-excluded tumors.

**SIGNIFICANCE:** The cellular and molecular programs driving T cell marginalization in solid tumors remain unclear. Here, we describe two CAF populations associated with T cell exclusion in human lung tumors. We demonstrate the importance of pairing molecular and spatial analysis of the tumor microenvironment, a prerequisite to develop new strategies targeting T cell-excluding CAF.

## INTRODUCTION

Lung cancer is the leading cause of cancer-related deaths worldwide, accounting for roughly 1.6 million deaths per year, with non-small cell lung carcinoma (NSCLC) being the most prevalent form (1). The partial success of immune checkpoint blockade in only a subset of NSCLC patients underscores the need for a better understanding of the determinants controlling anti-tumor immunity (2). In addition to high tumor mutational burden and PD-L1 expression levels in the tumor, CD8^+^ T cell density has been shown as a predictor of immunotherapy response (3, 4). By analyzing the T cell localization within the tumor, recent studies have revealed the importance of T cell infiltration into the tumor nests relative to the surrounding stroma (3,5,6). Understanding the mechanisms regulating T cell exclusion are therefore crucial to improve T cell-based therapies and patient outcomes.

Using real-time imaging of T cell dynamics in human NSCLC, we previously found that dense fibers oriented parallel to the tumor-stroma interface form a barrier around the tumor mass and limit T cell contact with tumor cells (7). However, the cellular sources and their extracellular matrix (ECM) programs remain unknown. Fibroblasts are known to shape lymphocyte compartmentalization in secondary lymphoid organs, where they produce distinct sets of chemokines and a complex ECM conduit system that serves as a scaffold along which dendritic cells and lymphocytes migrate and engage (8–10). While the role of fibroblasts in restricting immune cell localization is well established in spleen and lymph nodes, only recently has the tumor stroma emerged as a player in regulating local immune responses (11–14).

Given the growing evidence indicating that cancer-associated fibroblasts (CAF) can regulate tumor immunity and progression(11–14), CAF are becoming an important target for cancer treatment. TGFβ blockade and NOX4 inhibition were shown to act on CAF and facilitate T cell infiltration, leading to better responses to anti-PD-1/PD-L1 treatment in murine cancer models (6,15,16). Yet modulating and depleting CAF have led to opposite results in other tumor systems (16, 17) and has not yet managed to achieve clinical benefit in human cancer (18, 19). How to manipulate fibroblast properties for therapeutic purpose remains challenging, largely due to our limited understanding of the tumor CAF compartment and the mechanisms by which distinct CAF populations modulate anti-tumor immunity, including immune cell spatial organization.

The initial characterization of functional heterogeneity of CAF included description of inflammatory CAF (iCAF) and myofibroblastic CAF (myCAF) in mouse models of pancreatic cancer(20). Transcriptional signatures of these distinct CAF phenotypes have subsequently been found in human pancreatic and breast cancer(21, 22), as well as an additional subset, antigen-presenting CAF (apCAF)(22). iCAF are described as being found distal from the tumor site with a secretory phenotype whereas myCAF are characterized by activation and contractility genes and their close proximity to tumor cells(20). Prior studies have used single-cell RNA sequencing (scRNAseq) to profile CAF in various human cancers, including NSCLC(23), bladder(24), pancreas(22, 25), breast(21), head and neck(26), and liver(27). While the diversity of CAF is increasingly appreciated, the molecular programs of human fibroblast subsets and their discrete functional contributions to the tumor organization and T cell compartmentalization have not been resolved.

We reasoned that pairing scRNAseq profiling with high resolution spatial mapping would enable unbiased identification of CAF transcriptional subsets and uncovering their spatial organization in the context of the tumor microenvironment. Our scRNAseq analysis on 15 surgically resected NSCLC samples along with 12 paired adjacent tissue samples identified novel CAF subpopulations which we validated by profiling 35 tumors by multiplexed immunohistochemistry (IHC) (28). We analyzed the spatial organization of the stromal and immune cell populations, and revealed two CAF subsets with distinct ECM programs that were associated with CD3^+^ and CD8^+^ T cell exclusion from the tumor nests. Importantly, by applying high-resolution histological profiling on a large NSCLC cohort, our study characterizes both the intra-tumor and inter-tumor CAF and T cell heterogeneity.

## RESULTS

### Paired scRNAseq and IHC analysis identifies four CAF populations with distinct transcriptional profiles and structural organization in human NSCLC

To characterize the stromal cell compartment in NSCLC in an unbiased way, we profiled non-immune, non-tumor/epithelial cells isolated from 15 NSCLC samples and 12 paired adjacent tissue samples using the 10x Genomics scRNAseq platform (Figure 1A, Table 1). Using flow cytometry and mass cytometry by Time of Flight (CyTOF), we optimized the digestion and sorting protocols to maximize the stromal cell recovery while preserving cell integrity (Methods, Figures S1A, S1B). 33,742 cells were sequenced in total which contained 31,402 stromal cells after excluding contaminating immune cells, epithelial cells, and cells not passing quality control (Table 2). Using an unsupervised clustering, which integrates samples over different conditions and patients while modeling background noise (29, 30), we identified 28 clusters, including; 24 stromal cell clusters of variable abundance shared among samples (Figure S1C, table 2) as well as 4 clusters, either containing contaminating immune cells or cells with high mitochondrial content, that were excluded from future analysis. mRNA counts (unique molecular identifiers, UMI) per cluster and mitochondrial content per sample were similar (Figures S1C, S1D).

**Fig. 1.**
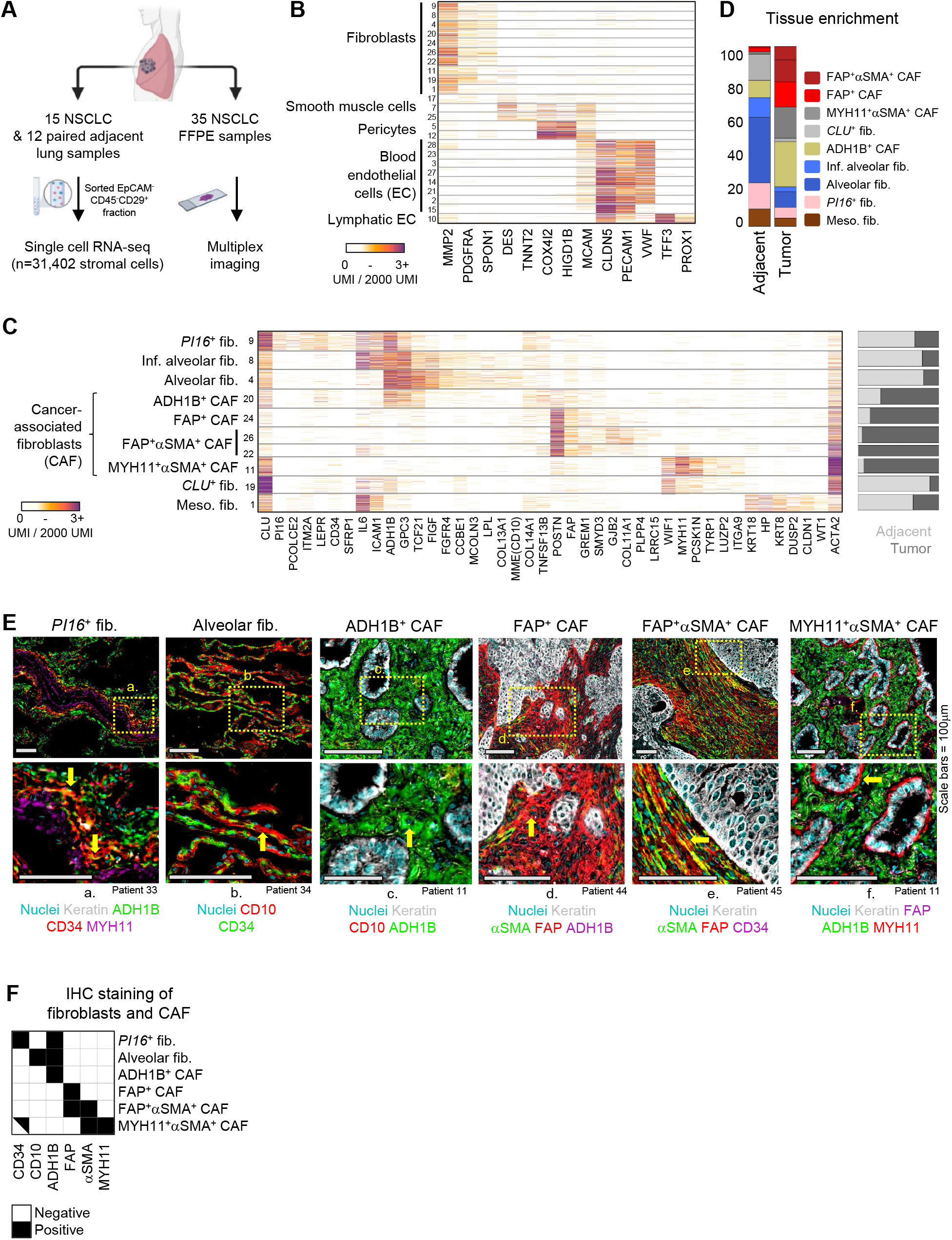
Paired scRNAseq and IHC analysis identifies four CAF populations with distinct transcriptional profiles and structural organization in human NSCLC. A, Tissue processing workflow for scRNAseq and IHC imaging of FFPE samples. B, scRNAseq mRNA counts (unique molecular identifiers, UMI) per cell (rows) of select stromal lineage marker genes (columns). Fibroblast, smooth muscle, pericyte, blood and lymphatic endothelial cell clusters are identified based on expression of marker genes such as, PDGFRA, DES, COX4I2, PECAM1, and TFF3, respectively. All cells displayed in this figure, and all subsequent similar scRNAseq figures, were downsampled to 2000 UMI. C, Extended gene lists highlighting gene expression profiles between the fibroblast subsets along with differing propensities for enrichment (right bar plot) in tumor (dark gray) or adjacent tissue (light gray). D, Averaged fibroblast composition in adjacent and tumor samples across all patients. The bar graph depicts the percentage of cells from each fibroblast subset among all fibroblasts. E, FFPE NSCLC sections were stained for fibroblast markers identified in scRNAseq results. All the scRNAseq-based fibroblast clusters (D) were detected utilizing IHC except meso. fib. and CLU^+^ fib., which were not in the scope of this study. Arrows highlight cells of interest (*PI16*^+^ fib.: CD34^+^ADH1B^+^MYH11^neg^, Alv. fib.: CD10^+^CD34^neg^, ADH1B^+^ CAF: ADH1B^+^CD10^neg^, FAP^+^ CAF: FAP^+^ADH1B^neg^αSMA^neg^; FAP^+^αSMA^+^ CAF: FAP^+^αSMA^+^CD34^neg^; MYH11^+^αSMA^+^ CAF: MYH11^+^FAP^neg^ADH1Β^neg^). See Figure S3 for other stainings. All scale bars are 100μm. F, IHC staining presentation for the main identified fibroblast and CAF clusters.

To unbiasedly dissect cell identities, we analyzed the mRNA counts of variably expressed genes across the 24 stromal cell clusters (Figure S1E). The cell clusters represented 3 major stromal cell compartments and expressed well reported lineage markers: fibroblasts (PDGFRA^+^, MMP2^+^), endothelial cells [EC, including both blood (CLDN5^+^, PECAM1^+^) and lymphatic (TFF3^+^, PROX1^+^) EC], and perivascular cells [PvC, including pericytes (MCAM^+^, COX4I2^+^), and smooth muscle (SM) cells (MCAM^+^, DES^+^)] (Figures 1B, S1F). Blood EC clusters included arteries, venules, tip cells, as well as two lung capillary subsets recently described as aerocytes and general capillaries(31) (Figure S2A). The PvC clusters enriched in tumor lesions included tumor pericytes, which expressed high amounts of RGS5 and multiple collagens (COL1A1, COL3A1, COL6A3), and a cluster expressing multiple immunomodulatory genes including CCL19 and CCL21 (Figure S2B). To be noted, IHC showed that the MCAM^+^ cells were restricted to vascular areas and were not found in the rest of the stroma (Figure S2C).

Further dissection of fibroblast populations identified multiple subsets with distinct transcriptional profiles and uneven abundances in the tumor lesion or the adjacent tissue (Figures 1C, 1D, Table 3). Based on this scRNAseq analysis, we identified genes associated with each cluster and defined antibody panels (Table 4) that enabled further characterization by multiplexed IHC (Figures 1E, 1F, S3A-B). Two clusters enriched in the adjacent lung tissue were characterized by co-expression of *MME* (CD10), *FIGF* (VEGFD), *FGFR4* (Figure 1C) and were annotated as alveolar fibroblasts (alv. fib.) based on their specific localization to the lung alveoli by IHC (Figures 1E, S3A). Interestingly, one of these clusters expressed high levels of inflammatory genes, including *IL6* and *ICAM1*, and was thus referred to as inflamed alveolar fibroblasts (inf. alv. fib.) (Figure 1C). Another cluster enriched in the adjacent lung was annotated as *PI16^+^* fibroblasts based on its co-expression of *PI16*, *CD34 and LEPR* (leptin receptor), localization to the blood vessel adventitia, and similarity to the universal *PI16^+^* fibroblasts described in Buechler et al., 2021 (Figures 1C, 1E, S3A) (32–34). The last adjacent tissue cluster, CLU^+^ fib., was characterized by high expression of *CLU* (clusterin) (Figure 1C).

Fibroblast clusters enriched in tumor samples were annotated as CAF. One CAF cluster displayed an expression profile similar to that of alv. fib., including expression of the broad adjacent tissue fibroblast marker, *ADH1B* (alcohol dehydrogenase 1B), and lower expression of the canonical CAF marker, *FAP*, and was referred to as ADH1B^+^ CAF (Figures 1C, 1E, S3A). ADH1B^+^ CAF could be distinguished from alv. fib. in IHC by their lack of CD10 expression and localization in the tumor lesion (Figures 1C, 1E, S3A, S3C). Three clusters showed strong expression of canonical activated CAF markers, *FAP*, *POSTN, LRRC15* and *GREM1* (23, 25) and were denoted as FAP^+^ CAF (Figures 1C, 1E, S3B). Another common CAF marker, *ACTA2* (αSMA) (35), was differentially expressed among the FAP^+^ CAF (Figures 1C, 1E, S3B) and clusters with high *ACTA2* expression were designated as FAP^+^αSMA^+^ CAF. Notably, a cluster that was highly enriched in a single patient (Table 2) shared both fibroblast genes (*PDGFRA*, *MMP2*, *COL1A1*, *BGN*) and mesothelial cell genes, such as keratins and WT1, and was therefore designated as mesothelial-like fibroblasts (meso. fib.) (36) (Figure 1C). An additional CAF cluster, MYH11^+^αSMA^+^ CAF, clearly distinct from the other CAF subsets, was characterized by the expression of *MYH11* (myosin heavy chain 11), *ACTA2* and intermediate levels of *CD34*, while lacking *ADH1B* and *FAP* expression (Figure 1C). Histological analysis of matched formalin-fixed paraffin-embedded (FFPE) tumor samples revealed a MYH11^+^αSMA^+^CD34^+^ADH1B^neg^FAP^neg^ cell population observed as a single layer of elongated CAF encapsulating tumor nests, in contrast to ADH1B^+^ CAF and FAP^+^ CAF that are spread throughout the stroma (Figures 1E, 1F, S3A). CyTOF confirmed the presence of the main fibroblast subsets identified through scRNAseq, including alv. fib., PI16^+^ fib., MYH11^+^αSMA^+^ CAF and FAP^+^αSMA^+^ CAF (Figure 2A).

**Fig. 2.**
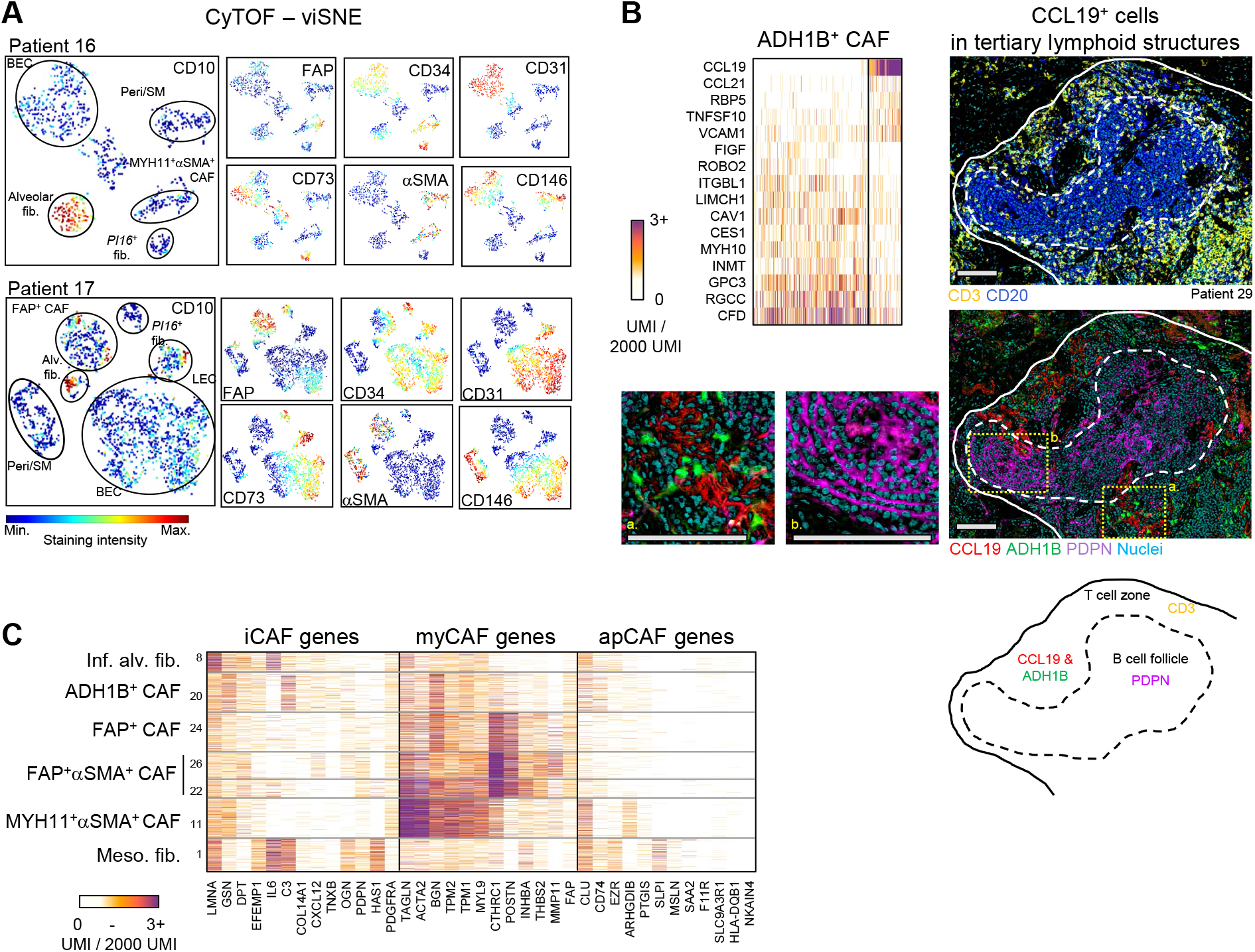
Further characterization of CAF subsets in human NSCLC. **A**, Stromal cell populations visualized with viSNE in CyTOF. EC, PvC and multiple fibroblast subsets can be distinguished with relatively few markers (CD10, CD31, CD34, CD73, FAP, CD146 and αSMA) **B**, (upper left panel) Highlighting CCL19 expressing cells within ADH1B^+^ CAF. These cells expressed high amounts of CCL19, CCL21, and VCAM1 and low levels of certain ADH1B^+^ CAF genes such as MYH10 and GPC3. (bottom and right panels) Multiplex IHC of a representative tertiary lymphoid structure. Podoplanin (PDPN) and CD20 marks follicular dendritic cells and B cells, respectively, in the B cell follicle, while the T cell zone is identified with CD3 staining. CCL19 and ADH1B staining show ADH1B^+^ fibroblasts surrounded by the secreted chemokine CCL19, specifically in the T cell zone. All scale bars are 100μm. **C**, myCAF, iCAF and apCAF gene signatures (Elyada et al., 2019; Öhlund et al., 2017) projected onto NSCLC CAF clusters.

Further analysis of ADH1B^+^ CAF revealed a subset of cells that expressed high levels of T cell-attracting and T cell-retention genes *CCL19*, *CCL21*, and *VCAM1*, reminiscent of fibroblastic reticular cells present in secondary lymphoid organs (37) (Figure 2B). IHC staining of CCL19 confirmed the specific localization of these fibroblasts to tertiary lymphoid structures, with clear preferential enrichment for the T cell zone (Figure 2B). In some cases, the B cell zone was delineated by podoplanin expression, which marks follicular dendritic cells that were not captured by scRNAseq, likely due to their low abundance (Figure 2B).

Taken together, our combined IHC and single-cell analysis has defined diverse fibroblast populations with distinct molecular and spatial patterns in human NSCLC. By enriching for stromal cells from a large NSCLC cohort, we achieved highly granular scRNAseq characterization and uncovered CAF populations undescribed to date, including a single layer of MYH11^+^αSMA^+^ CAF bordering tumor cells in a fraction of NSCLC lesions. The four CAF subsets described here expand upon the iCAF, myCAF, and apCAF profiles described in pancreatic tumors(20, 22). Our analysis suggests that in human lung tumors, myCAF include both FAP^+^ CAF, FAP^+^αSMA^+^ CAF and MYH11^+^αSMA^+^ CAF highlighting the transcriptomic and spatial complexity of this population (Figure 2C).

### ADH1B^+^ CAF and FAP^+^ CAF stratify NSCLC into two main stromal patterns associated with tumor stage and histology

Analysis of the fibroblast composition as determined by the scRNAseq indicated that low stage tumors were dominated by ADH1B^+^ CAF with or without MYH11^+^αSMA^+^ CAF, while higher stage tumors were enriched for FAP^+^ CAF and FAP^+^αSMA^+^ CAF (Figures 3A, 3B, S4A). To test the dichotomy between ADH1B^+^ and FAP^+^ CAF enrichment, we leveraged a larger cohort of 35-patient FFPE samples and quantified the tumor area covered by ADH1B and FAP using IHC. This unbiased analysis showed that the stroma of NSCLC is significantly dominated for either ADH1B^+^ or FAP^+^ CAF (hypergeometric test, *p = 0.008*) (Table 5, Figure 3C). MYH11^+^αSMA^+^ CAF were observed in half of ADH1B^+^ CAF rich samples (9/18) (Figure 3C) but they were not observed in FAP^+^ CAF rich samples, corroborating the scRNAseq analysis that showed a correlation between ADH1B^+^ CAF and MYH11^+^αSMA^+^ CAF (Figure 3A). FAP^+^ CAF rich samples showed highly variable expression of αSMA (Figure 3B), in line with the variable *ACTA2* expression seen across the scRNAseq FAP^+^ CAF clusters (Figure 1C).

**Fig. 3.**
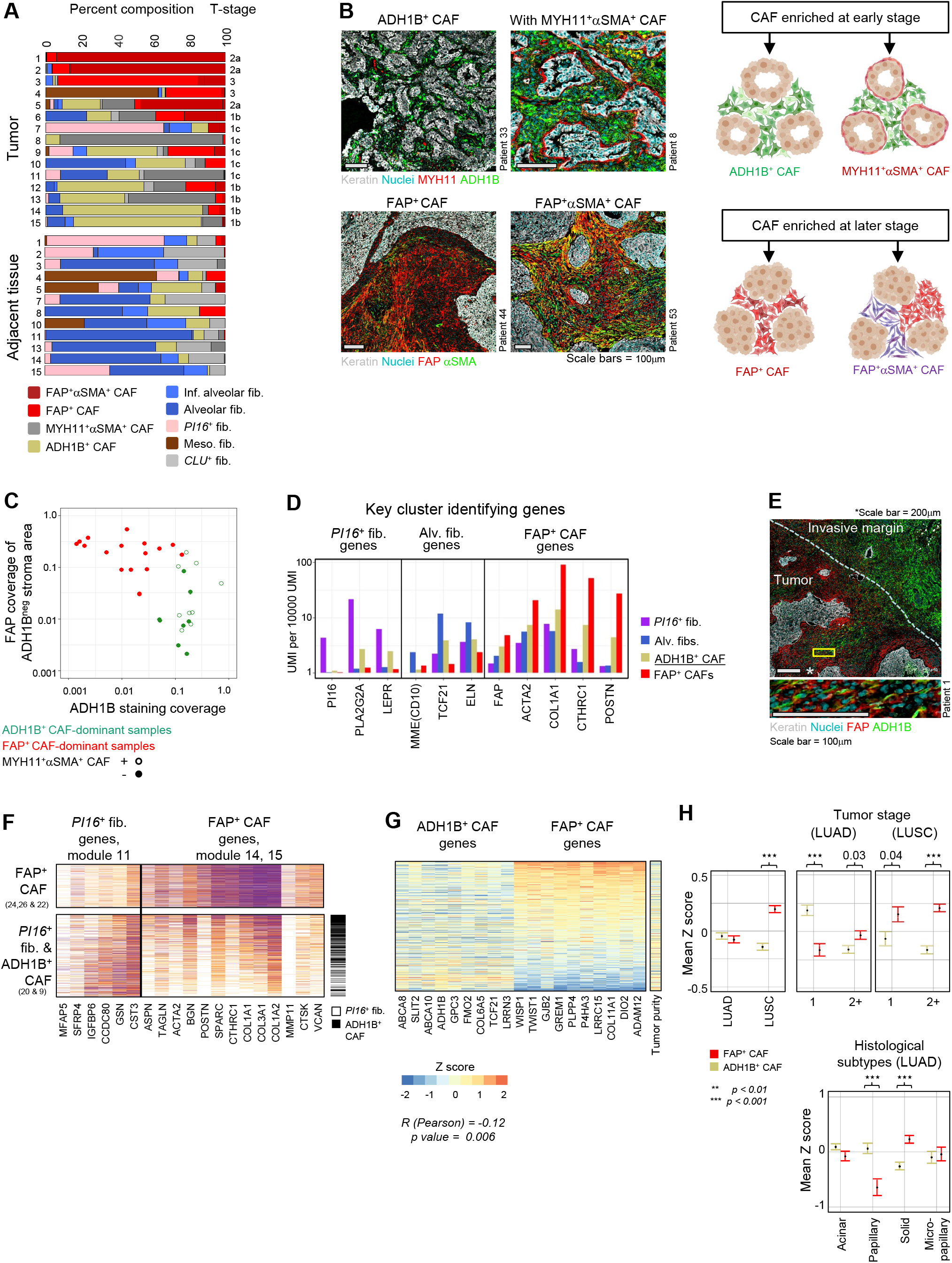
ADH1B^+^ CAF and FAP^+^ CAF stratify NSCLC into two main stromal patterns associated with tumor stage and histology. **A**, Fibroblast subset composition, displayed by percentages, in individual tumor and adjacent tissue samples from the 15 scRNAseq patients. **B**, (Top left panels) ADH1B^+^ CAF rich patients showing ADH1B presence throughout the stroma. ADH1B^+^ CAF rich patients may present with (bottom left panel) or without (top left panel) a distinct single cell layer of MYH11^+^αSMA^+^ CAF at the tumor border. (Bottom left panels) FAP^+^ CAF rich patients with FAP staining throughout the stroma. The patients shown demonstrate the variable αSMA presentation in FAP^+^ cells. All scale bars are 100μm. (Right panels) Cartoon illustrating the observed presentation of multiple CAF subsets in NSCLC. **C**, ADH1B and FAP staining in the IHC cohort. ADH1B staining coverage in the stroma is shown on the X axis. FAP staining coverage in the stroma on regions that did not stain for ADH1B are shown on the Y axis. Tumors show significant preference for either ADH1B or FAP, with less than 5% coverage of the opposing stain. The 5% cutoff was selected after performing hypergeometric tests for 10 thresholds, at 5% increments, between 5% and 50%. The Bonferroni correction adjusted p value is 0.008. **D,** Mean expression of selected genes highlighting ADH1B^+^ CAF intermediate expression of *PI16*^+^ fib., alv. fib., and FAP^+^ CAF-associated genes. **E**, Tumor sample with an extensive invasive margin that displays a spectrum of ADH1B to FAP staining. (Zoom, bottom panel) Cells appearing to transition from ADH1B to FAP expression. Top panel scale bar is 200μm, bottom panel scale bar is 100μm. **F**, Expression of *PI16*^+^ fib. and FAP^+^ CAF module genes in *PI16*^+^ fib., ADH1B^+^ CAF, and FAP^+^ CAF. Based on gene expression patterns, ADH1B^+^ CAF appear to occupy an intermediate state of activation between *PI16*^+^ fib. and FAP^+^ CAF. **G**, Relative expression, displayed by Z score, of ADH1B^+^ CAF and FAP^+^ CAF-associated genes in TCGA LUAD bulk-RNAseq samples. ADH1B^+^ CAF and FAP^+^ CAF genes are significantly anticorrelated (Pearson) *R = -0.12* and *p = 0.006*. The sample tumor nuclei count is used as a proxy of tumor purity and shows a relatively even distribution. **H,** TCGA LUAD mean Z score and standard error of mean (SEM) of ADH1B^+^ CAF and FAP^+^ CAF gene signatures stratified by tumor subtype (left and middle panels) or stage (right panel). Z score calculation is listed in methods and significance is calculated by independent t test (right panel).

To study the transcriptional programs behind ADH1B^+^ CAF and FAP^+^ CAF and to better understand their relationship to adjacent tissue fibroblasts, we analyzed gene expression covariance patterns across ADH1B^+^ CAF, FAP^+^ CAF, alv. fib. and *PI16*^+^ fib. We identified groups of co-expressed genes (gene modules) with distinct expression patterns across these fibroblast populations (Figure S4B). FAP^+^ CAF upregulated activation genes (modules 14, 15) including multiple collagen genes (*COL1A1*, *COL3A1*) that contribute to tissue stiffness(38) and other ECM genes such as biglycan (*BGN*) that can promote tissue mineralization (39) (Figure S4B). FAP^+^ CAF expressed low levels of the alv. fib. genes, including the fibroblast transcription factor *TCF21* (40), the marker *MME*(CD10) as well as the ECM gene elastin (*ELN*) which is critical for normal lung physiology(41) (Figure 2D). ADH1B^+^ CAF expressed intermediate levels of FAP^+^ CAF activation genes (Figure 3D), and a subset of samples showed a gradient of ADH1B^+^ CAF to FAP^+^ CAF from the invasive margin to the tumor center (Figure 3E) with some cells co-expressing both markers, suggesting that ADH1B^+^ CAF represent a range of lowly activated fibroblasts. ADH1B^+^ CAF also shared genes with both *PI16*^+^ fib. and alv. fib., which may point towards both lung fibroblast types as their potential cellular sources (Figures 3D, S4C). Interestingly, the scRNAseq data showed that ADH1B CAF cells express a gradient of FAP^+^ CAF and *PI16*^+^ fib. genes from cells with high expression of *PI16* genes and low FAP genes to cells with low *PI16* genes and high FAP genes. This further supports the hypothesis that ADH1B^+^ CAF are a lowly activated form of fibroblast and may derive from *PI16*^+^ fib. (Figure 3F).

Leveraging our scRNAseq datasets, we created gene signatures for ADH1B^+^ CAF and FAP^+^ CAF by selecting for genes with highly specific expression in their corresponding CAF populations in contrast with all other cell types. With these signatures we scored the Cancer Genome Atlas (TCGA) lung adenocarcinoma (LUAD) samples by their expression of ADH1B^+^ CAF and FAP^+^ CAF genes and revealed an anticorrelation between the two scores (*p = 0.006*) (Figures 3G, S4D, Table 6), supporting the two distinct CAF profiles observed across NSCLC patients in scRNAseq and histology (Figures 3A-C) (42). Analysis of tumor purity, estimated by tumor nuclei abundance, did not reveal clear association with ADH1B^+^ CAF or FAP^+^ CAF genes (Figure 3G), confirming that contaminating adjacent tissue was not a major contributor to the ADH1B^+^ CAF signal. Further analysis of TCGA data showed that ADH1B^+^ CAF genes were significantly increased in stage 1 tumors, LUAD and the papillary LUAD subtype, whereas FAP^+^ CAF were enriched in later stage tumors, LUSC, and the LUAD solid subtype (Figure 3H). LUAD across tumor stages confirmed our observation that ADH1B^+^ CAF and FAP^+^ CAF were correlated with lower and higher stage tumors, respectively. Similar associations were observed in our in-house FFPE cohort (Table 1). Altogether, we showed that ADH1B^+^ and FAP^+^ CAF phenotypes were correlated with tumor stage and clinically relevant histological subtypes (42, 43), suggesting that molecular characterization of fibroblasts could refine clinical categorization of NSCLC tumors.

### ADH1B^+^ CAF and FAP^+^ CAF correlate with immune cell composition and not with T cell localization

Given the data showing that CAF contribute to regulating tumor immunity (11,12,44) we investigated the different ligands expressed by CAF populations found in human NSCLC (Figure 4A). Increased expression of the cytokines *IL34* and *CSF1* suggested macrophage regulation (45) by ADH1B^+^ CAF, whereas FAP^+^ CAF might attract eosinophils/basophils via *CCL11* (46), as well as CCR5^+^ T cells and monocytes through *CCL3* and *CCL5* chemokines (47–49). Notably, the high levels of *CCL21* and *TNFSF13B* (BAFF) in ADH1B^+^ CAF mainly come from the CCL19-expressing ADH1B^+^ cells specifically found in TLS (Figures 4A, 2B), likely contributing to naïve T cell attraction and B cell survival in these structures (37). MYH11^+^αSMA^+^ CAF expressed increased levels of TSLP (thymic stromal lymphopoietin) which can stimulate the maturation of immune cells that express both *IL7R* and *CRLF2* genes forming the heterodimeric TSLP receptor, such as certain dendritic cells (50). To further investigate the contribution of different CAF populations to shaping the immune microenvironment, we used multiplex imaging on FFPE tissue to histologically profile the CAF subset composition of a large cohort of NSCLC samples that we had previously studied using scRNAseq of purified immune cells(29). We demonstrated a significant association (Pearson, *R=0.62, p = 0.01*) between the presence of FAP^+^ CAF and the enrichment of inflammatory *SPP1*^+^ monocyte-derived macrophages, IgG^+^ plasma cells, and PD1^+^ T cells (Figure 4B). These immune cell types were recently described as part of a cellular module termed Lung Cancer Activation Module (LCAM)(29). We then validated this CAF-immune association in the TCGA LUAD cohort. There was a significant correlation between CAF phenotype and the LCAM score (*R = 0.66*, *p < 1e^-10^*), supporting that FAP^+^ CAF rich samples are linked to more inflammatory and activated immune cells, LCAM^high^, in LUAD (Figure 4C).

**Fig. 4.**
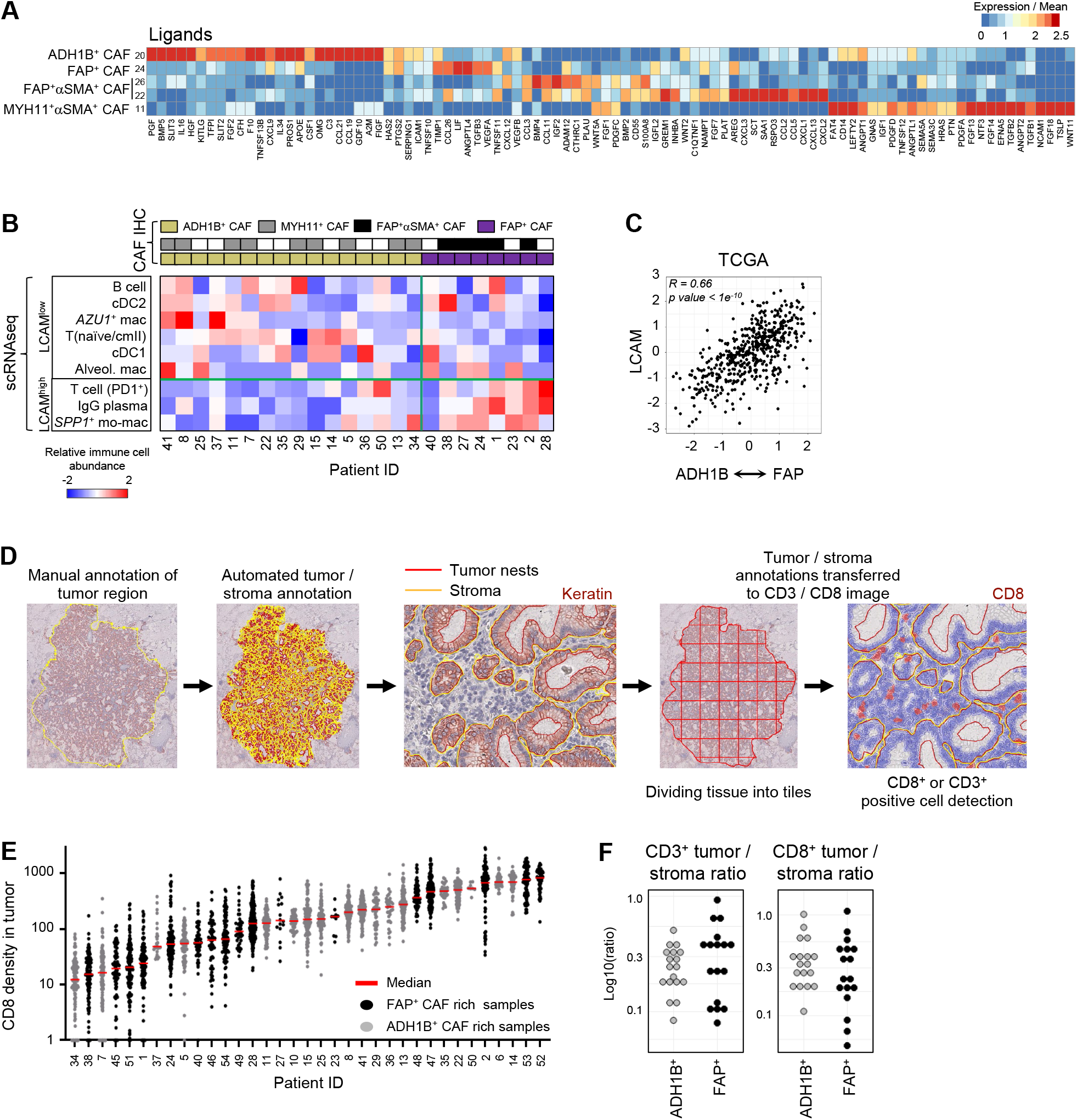
ADH1B^+^ CAF and FAP^+^ CAF correlate with immune cell composition and not with T cell localization. **A**, Gene expression over mean of highly variable immunomodulatory ligands in CAF clusters. **B**, Immune composition of scRNAseq tumor samples from (Leader et al., 2021). CAF phenotype is identified by IHC on the matched FFPE samples and then used to stratify samples. The relative abundance of each cell population within its respective compartment, i.e.; PD1^+^ T cells amongst all T cells, is calculated and then scaled across all tumors for the respective Z score value. LCAM score is significantly correlated with CAF phenotype (Pearson) *R = 0.62*, *p = 0.01*. **C**, Estimating the correlation between CAF phenotype and LCAM in TCGA LUAD samples. Each patients’ mean ADH1B^+^ CAF gene signature is subtracted from their mean FAP^+^ CAF gene signature and the resulting values are correlated with estimate LCAM score. The corresponding Pearson correlation values are shown. **D**, Schematic of QuPath methodology for tiling and T cell quantification. **E**, CD8^+^ cell infiltration into tumor nests in each patient (columns). Each point represents an individual 1000μm x 1000μm tile (all other tiling is 500μm x 500μm). **F**, IHC quantification of CD3^+^ or CD8^+^ cells per mm^2^ in the tumor and the tumor / stroma CD3^+^ or CD8^+^ cells per mm^2^ ratio. Tumor samples are stratified by their stroma profile (ADH1B^+^ CAF rich or FAP^+^ CAF rich) and no significant difference was observed.

Given that the spatial distribution of T cells is a predictor of clinical response to immune checkpoint blockade (51), we used an unbiased cell quantification method to measure T cell infiltration in the tumor nests and identified a wide range of infiltration levels across the cohort (Figures 4D-F). Importantly, there was no observed association between any of ADH1B^+^ and FAP^+^ CAF-rich profiles and CD3^+^ and CD8^+^ T cell localization (Figures 4E-F, Table 7). Taken together, ADH1B^+^ CAF and FAP^+^ CAF stratify tumor lesions by two levels of fibroblast activation and correlate with the immune phenotype, but not with T cell spatial distribution.

### MYH11^+^αSMA^+^ CAF are correlated with decreased T cell infiltration in tumor nests

The lack of correlation between ADH1B^+^ CAF or FAP^+^ CAF with CD3^+^ or CD8^+^ T cell infiltration contrasts with the general idea that activated fibroblasts orchestrate T cell exclusion, raising the hypothesis that fibroblast subsets other than ADH1B^+^ or FAP^+^ CAF could be involved. To investigate if MYH11^+^αSMA^+^ CAF, which form a single cell layer around tumor nests in a fraction of early-stage tumors, could also impact T cell tumor infiltration, we subdivided stage 1 patients based on the presence of MYH11^+^αSMA^+^ CAF at the tumor border (Figure 5A). In tumor lesions containing MYH11^+^αSMA^+^ CAF, the tumor-to-stroma ratio of infiltrating CD3^+^ or CD8^+^ cells was significantly lower (Figure 5B, right graphs), consistent with a decreased infiltrating CD3^+^ or CD8^+^ T cell density in the tumor (Figure 5B, left graphs) (Table 7). In addition, their high expression of *TGFB1* and *TGFB2* (Figure 4A) is in line with previous findings linking TGFβ and T cell exclusion (6, 52). These results suggested that MYH11^+^αSMA^+^ CAF, with their peri-tumoral location, may decrease T cells infiltration into tumor nests.

**Fig. 5.**
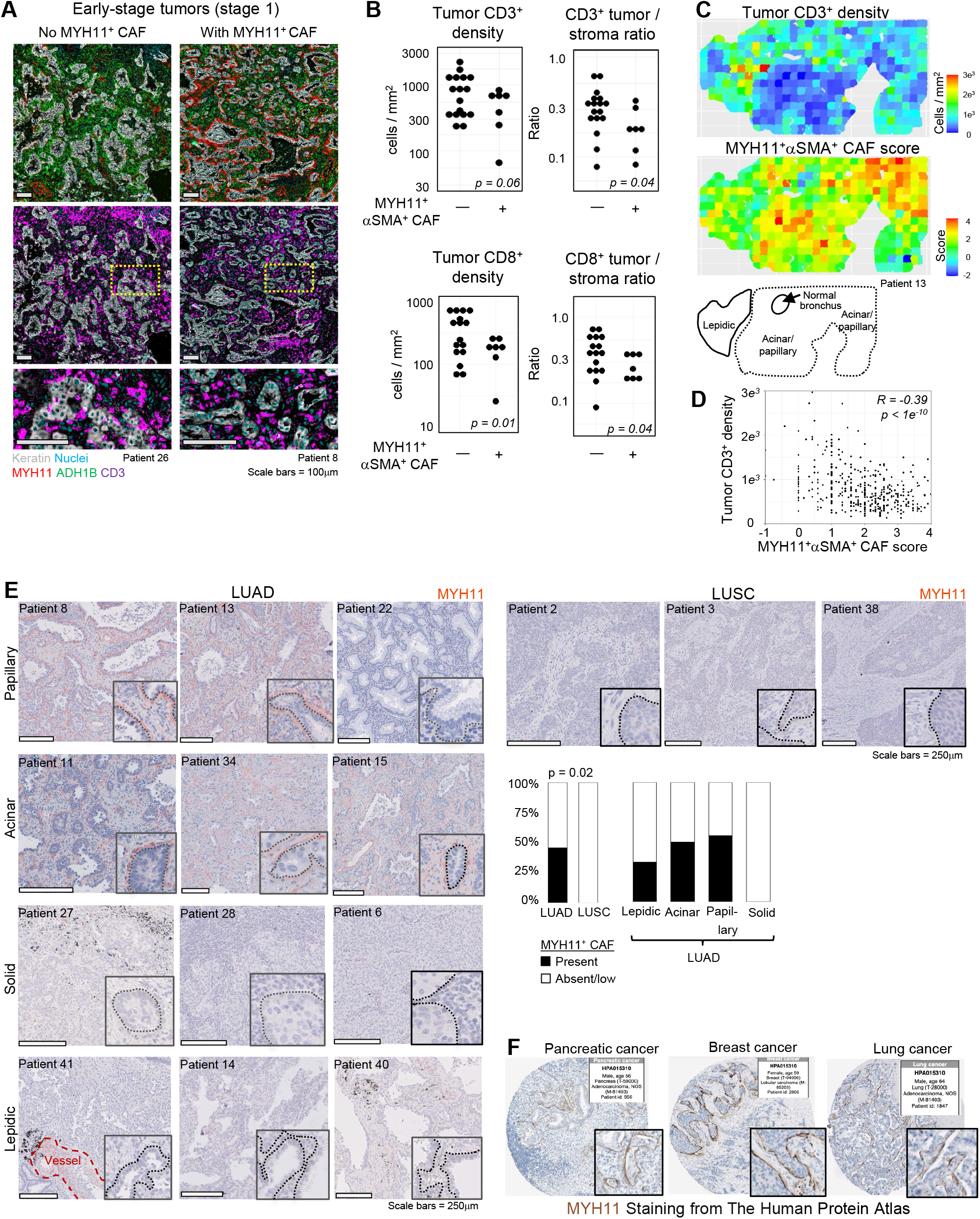
MYH11^+^αSMA^+^ CAF are correlated with decreased T cell infiltration in tumor nests. **A**, Representative examples of IHC stains from NSCLC tumors with and without MYH11^+^αSMA^+^ CAF present, showing CD3^+^ cell exclusion from tumor nests when MYH11^+^αSMA^+^ CAF are present. **B**, The presence or absence of MYH11^+^αSMA^+^ CAF demonstrates significant differences in tumor infiltrating CD3^+^ or CD8^+^ cells per mm^2^ (left) and the ratio of CD3^+^ or CD8^+^ cells per mm^2^ in the tumor versus stroma (right). Only early stage (tumor stage 1) patients were included to eliminate bias due to MYH11^+^αSMA^+^ CAF rarely being found at later stage. **C**, 500×500μm tiles of both MYH11^+^αSMA^+^ CAF score, estimating tumor proximity of MYH11^+^αSMA^+^ cells by quantifying their enrichment within 10μm from tumor cells versus regions 20μm-30μm from tumor cells, and tumor infiltrating CD3^+^ cells per mm^2^. (Bottom panel) Histological scoring of a tumor lesion highlighting that a high MYH11^+^αSMA^+^ CAF score is associated more with acinar/papillary phenotype, rather than lepidic. **D**, Quantification of results in F, a high MYH11^+^αSMA^+^ CAF score is significantly anti-correlated (Pearson) with the number of tumor infiltrating CD3^+^ cells relative to the stroma. **E**, Representative images of MYH11 staining in multiple pathologies and histological subtypes. All scale bars are 250μm. (Barplot) MYH11^+^αSMA^+^ CAF distribution in different pathologies and histological subtypes in NSCLC. **F**, MYH11 staining from The Human Protein Atlas.

Within our cohort, MYH11^+^αSMA^+^ CAF were found enriched in LUAD samples, especially in the acinar/papillary subtypes, while neither the solid subtype of LUAD, nor LUSC samples contained MYH11^+^αSMA^+^ CAF lining tumor nests (Figure 5E). Interestingly, the IHC image bank of the Human Protein Atlas showed that a similar peritumoral MYH11 staining pattern as one layer was observed in a fraction of samples of pancreatic and breast cancer (Figure 5F), suggesting that these CAF may be present in additional cancer types.

While most tumor lesions were characterized by either high or low MYH11^+^αSMA^+^ CAF presence, a fraction of tumors showed local heterogeneity. We assessed the intensity of CAF barrier at the tumor boundary in 500μm-by-500μm tiles using the abundance of MYH11^+^αSMA^+^ cells in the stroma close to (<10μm) versus distant from (20μm-30μm) tumor cells, which is referred to as the MYH11^+^αSMA^+^ CAF score (Figures 5C-D, S5). This automated analysis found that locations where MYH11^+^αSMA^+^ CAF were present had significantly lower tumor T cell density in two independent samples, highlighting that local spatial organization may be driving inter-tumor differences (Figure 5D). Additionally, histological analysis of the tumor lesion by a pathologist found that regions with high MYH11^+^αSMA^+^ CAF score were predominantly acinar/papillary, and lepidic regions at the tumor edge had a lower score (Figure 5C, bottom panel). Altogether, these results show that MYH11^+^αSMA^+^ CAF are a single layer of elongated cells associated with T cell marginalization both across NSCLC tumor samples and within tumor lesions.

### FAP^+^αSMA^+^ CAF define regions of poor T cell infiltration within tumor lesions and are coupled with dense ECM deposition

Spatial analysis of FAP^+^ CAF rich samples revealed that FAP^+^αSMA^+^ CAF, a subset of FAP^+^ CAF in scRNAseq, could also explain CD3^+^ and CD8^+^ T cell infiltration within tumors. We measured αSMA coverage and T cell density in the stroma in 500μm-by-500μm sections across the tumor lesion (Figure 6A) and revealed that regions dense in αSMA are poorly infiltrated by T cells (*R = -0.48*, *p = 1e^-10^*) (Figure 6B). This anticorrelation was replicated across different tumors (Figures S6A-C) and suggested that FAP^+^αSMA^+^ CAF directly restrict T cell motility. Notably, a high fraction of FAP^+^ CAF rich samples presented with several layers of FAP^+^αSMA^+^ CAF lining tumor nests that delineated regions devoid of T cells (Figure 6C), and this was corroborated by the finding that regions of high stromal αSMA coverage were associated with low T cell infiltration in the tumor nests (Figure S6C). In addition, inter-tumor αSMA heterogeneity showed a trend towards an anti-correlation with T cell infiltration in tumor nests (Figures S6D-E).

**Fig. 6.**
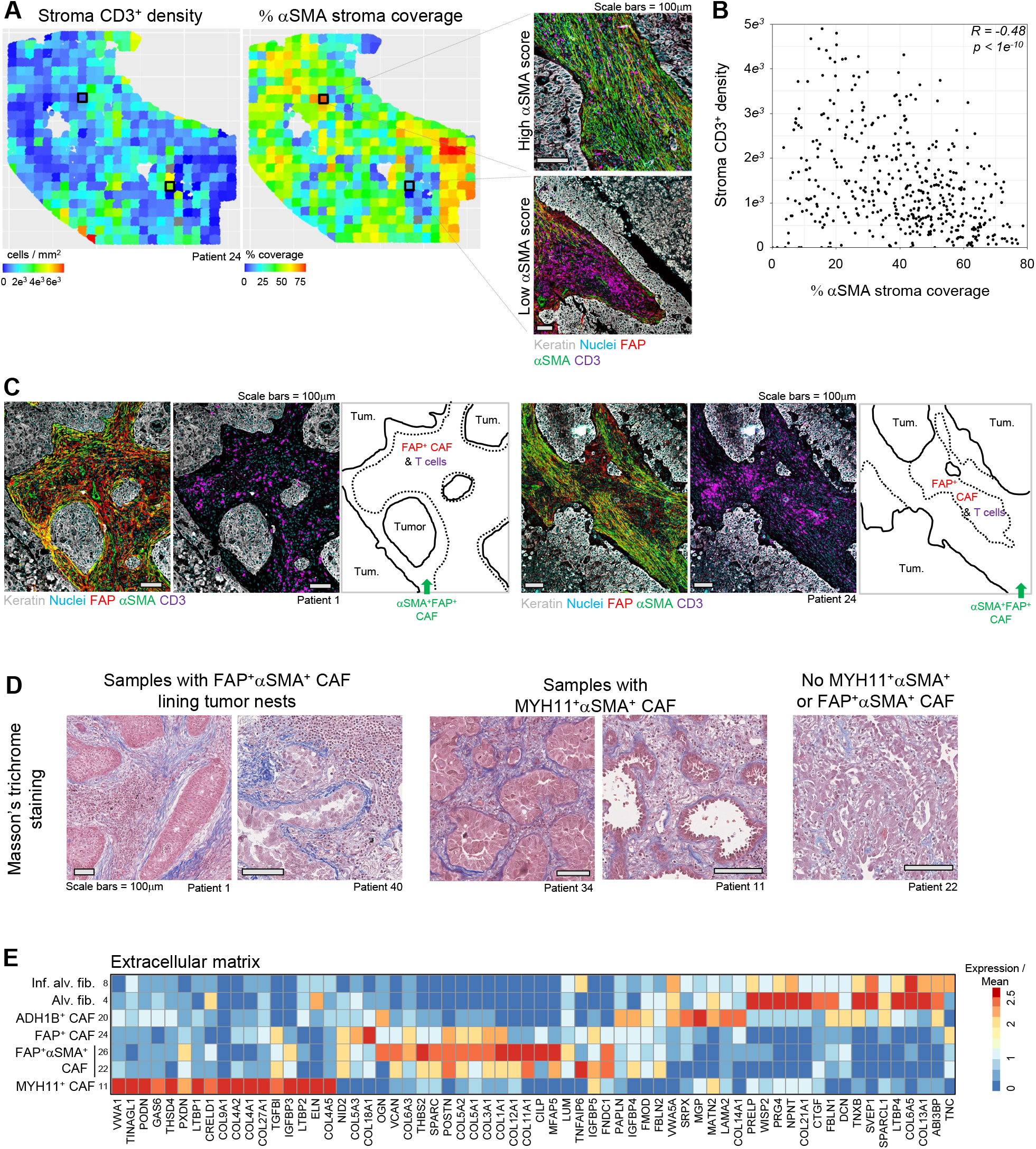
FAP^+^αSMA^+^ CAF define patterns of poor T cell infiltration within tumor lesions. **A**, (Left panel) Intra-tumoral heterogeneity of αSMA coverage (middle panel) and CD3^+^ cell density in the stroma in 500×500μm tiles. (Right panel) Representative examples of tiles showing regions with high or low levels of αSMA. **B**, Quantification of αSMA coverage and CD3^+^ density in each tile (points) as defined in A, showing a significant anticorrelation (Pearson) of αSMA coverage and CD3^+^ cell density. **C**, Dense αSMA staining at tumor border associates with decreased CD3^+^ cell abundance. The green arrow highlights border regions with high αSMA and low CD3^+^ cells. **D**, Masson’s trichrome stains highlighting increased ECM at the tumor boundary in samples containing MYH11^+^αSMA^+^ or FAP^+^αSMA^+^ CAF. **E**, Averaged gene expression of highly variable ECM genes in CAF clusters.

Based on prior studies showing that the ECM plays a role in T cell exclusion and immunosuppression(7,53,54), we postulated that FAP^+^ αSMA^+^ CAF may also express a specific ECM profile involved in regulating T cell localization. Masson’s trichrome staining revealed a high density of fiber deposition at the tumor border in both MYH11^+^αSMA^+^ and FAP^+^αSMA^+^ CAF-containing tumor lesions, suggesting that FAP^+^αSMA^+^ CAF are depositing a fibrillar barrier limiting T cell access to tumor cells (Figure 6D). Analysis of the scRNAseq showed that FAP^+^ αSMA^+^ CAF expressed high levels of the fibrillar collagen *COL11A1* (55), and *COL12A1* (Figure 6E). Notably, the ECM program of MYH11^+^αSMA^+^ CAF was distinct from FAP^+^αSMA^+^ CAF. MYH11^+^αSMA^+^ CAF expressed *COL9A1*, *COL27A1*, and a distinct type of sheet-forming, basement membrane collagens, *COL4A1* and *COL4A2*, which are found lining vessels and various epithelial layers(56, 57) (Figure 6E). Our data suggest that FAP^+^αSMA^+^ CAF and MYH11^+^αSMA^+^ CAF are two distinct types of fibroblasts forming the physical barrier of fibers which was previously observed around tumor nests and shown to restrict T cell infiltration into tumor islets in viable slices of human NSCLC lesions (7). Thus, the specific spatial distribution of MYH11^+^αSMA^+^ CAF and FAP^+^αSMA^+^ CAF and their unique ECM profiles may drive T cell exclusion in NSCLC and represent potential therapeutic targets, leading to a proposed model that refines the CAF landscape in human NSCLC (Figure 7).

**Fig. 7.**
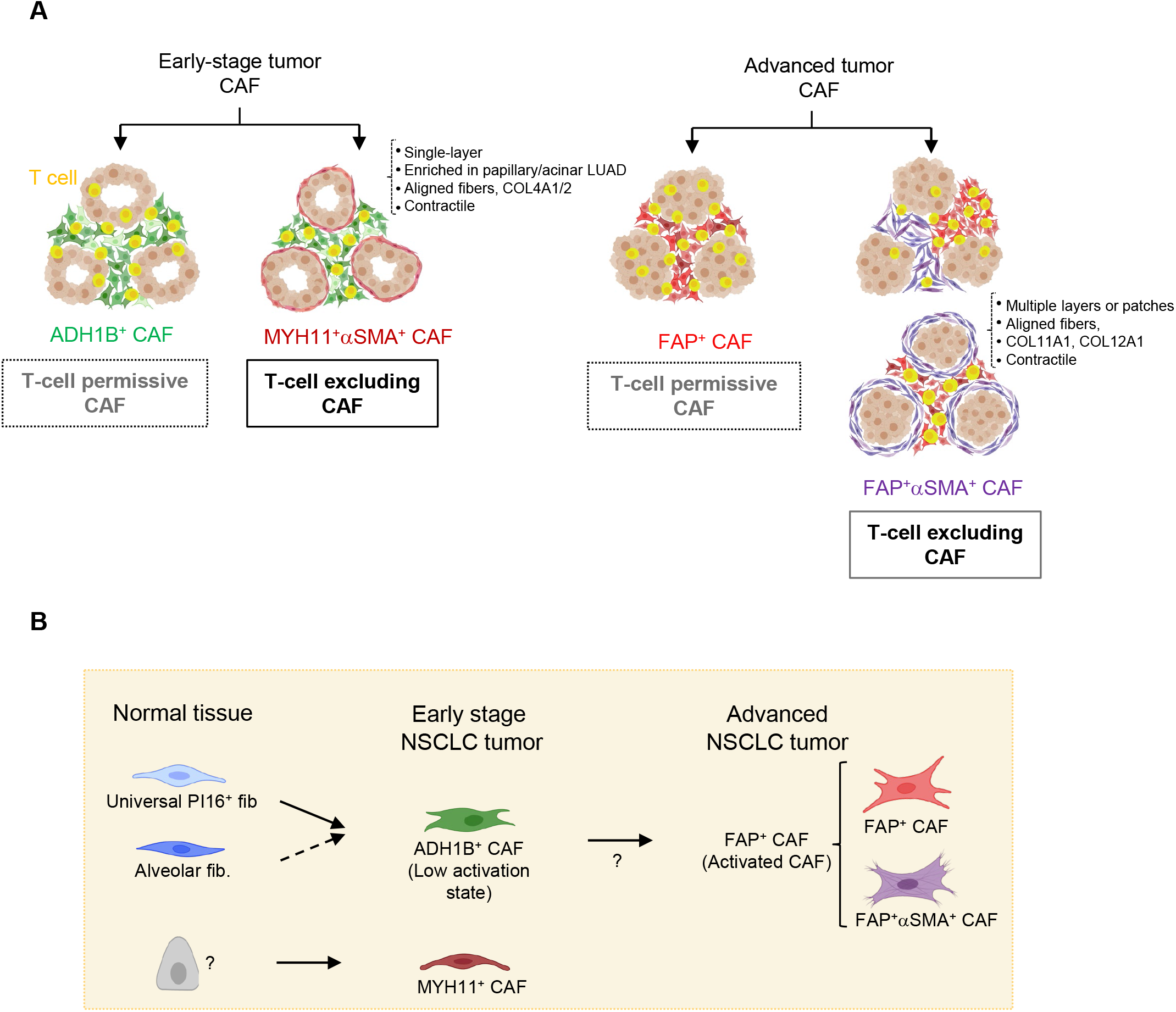
Working model. **A**, Graphical illustration of all stroma presentations found in this study. NSCLC samples enriched in ADH1B^+^ CAF throughout the stroma can be found with or without a single-cell layer of MYH11^+^αSMA^+^ CAF lining tumor cell aggregates. Those with MYH11^+^αSMA^+^ CAF show increased T cell exclusion from the tumor nests. NSCLC samples enriched in FAP^+^ CAF are found with variable abundance of FAP^+^αSMA^+^ CAF. Stromal regions with high αSMA have reduced T cell accumulation, and tumor nests surrounded by several layers of FAP^+^αSMA^+^ CAF show a lower T cell infiltration. **B**, Cartoon depicting the general distribution of fibroblast and CAF populations in adjacent lung tissue, early-stage NSCLC and advanced NSCLC, as well as the potential differentiation trajectories.

## DISCUSSION

The majority of patients fail to achieve clinical benefit using standard immune checkpoint blockade, and as such, novel combination approaches are required to improve response (58). Patients with T-cell excluded tumors have poor response to immune checkpoint blockade compared to those with T-cell infiltrated tumors(3,5,6) which raises the possibility that targeting the mechanism of T cell exclusion would improve clinical responses. To this end, our study provides a comprehensive map of the fibroblast compartment in human lung tumors at the single cell level and with spatial resolution. We define the molecular and functional diversity of the fibroblast compartment of lung tumors and determine how distinct CAF subsets may influence the immune cell composition as well as T cell spatial organization.

Our analysis shows that the stroma in NSCLC lesions is dominated by either lowly activated ADH1B^+^ CAF, with or without MYH11^+^αSMA^+^ CAF, or highly activated FAP^+^ CAF, with variable αSMA levels (Figure 7). ADH1B^+^ CAF have higher activation levels than fibroblasts found in the normal lung tissue, as seen by their expression profile and enrichment in the tumor lesion, as well as by the clear spatial distinction between CD10^+^ alv. fib. in adjacent tissue and ADH1B^+^CD10^neg^ CAF in the tumor as seen in multiplex IHC. FAP^+^ CAF, by contrast, show dramatic transcriptional differences from adjacent tissue fibroblasts, including high expression of many previously established CAF markers like *FAP*, *POSTN*, and *COL1A1* (23, 25). FAP^+^ CAF and FAP^+^αSMA^+^ CAF represent higher activation states compared to ADH1B^+^ CAF and occasionally form spatial gradients of ADH1B^+^-to-FAP^+^ CAF, suggesting that ADH1B^+^ CAF may contribute to the FAP^+^ CAF pool. Interestingly, ADH1B^+^ CAF express transcriptional programs of both alv. fib. and *PI16*^+^ fib., suggesting that these two lung tissue cell types could give rise to ADH1B^+^ CAF. *In vivo* fate-mapping experiments will be needed to further investigate this possibility.

We have also shown that ADH1B^+^ CAF and FAP^+^ CAF are associated with pathological and histological subtypes, and tumor stage, ADH1B^+^ CAF being associated with the adenocarcinoma papillary subtype and stage 1 whereas FAP^+^ CAF being enriched in the adenocarcinoma solid subtype and squamous cell carcinomas, and in later stage. This association between CAF populations and histological subtypes, which correlate with prognosis(43), may shed light on the molecular programs behind the different NSCLC subtypes, and inform clinical trial inclusion criteria when therapeutically targeting CAF subsets. Furthermore, our staining protocol for these CAF subsets may help refine the categorization of histological subtypes. Beyond subtype and stage, we found a significant association between FAP^+^ CAF and the LCAM inflammatory/activated immune phenotype (including *SPP1*^+^ Mo-Macs, IgG plasma cells, and PD1^+^ T cells) which we previously described (29). This observation, in conjunction with the distinct immunomodulatory profiles of ADH1B^+^ CAF and FAP^+^ CAF, suggests that CAF participate in shaping the immune response to the tumor.

Prior studies have suggested that activated CAF may play a role in T cell exclusion (6,15,16). Notably, FAP^+^ CAF do not correlate with the T cell distribution pattern in NSCLC, an important factor to keep in mind when developing therapeutics and may explain why strategies targeting FAP^+^ CAF have failed in human clinical trials so far (18, 59). In contrast, we have found two distinct CAF populations with specific molecular programs and spatial organizations that contribute to T cell exclusion. First, FAP^+^αSMA^+^ CAF are significantly correlated with regions of T cell exclusion in the tumor stroma and can form multiple layers at the tumor boundary and restrict T cell contact with tumor cells. On the other hand, in a fraction of adenocarcinomas MYH11^+^αSMA^+^ CAF form a single cell layer lining tumor nests, and are significantly correlated with immune cell exclusion from tumor regions, both within cancer lesions and across tumor samples.

FAP^+^αSMA^+^ CAF and MYH11^+^αSMA^+^ CAF correspond to two clearly distinct fibroblast subsets, as observed through the scRNAseq data and in line with their presence in distinct tumors. Notably, they display similarities, including high expression levels of ECM and contractility genes, which may imply that they influence T cell spatial distribution through similar mechanisms, with the production of fibers limiting T cell access to cancer cells. Supporting this, we previously showed that ECM fiber orientation and density dictate the motility of T cells within the stroma and restrict their interactions with tumor cells in human NSCLC (7). The profound differences in their matrix programs (including type IV and IX collagens for MYH11^+^αSMA^+^ CAF and type XI and XII collagens for FAP^+^αSMA^+^ CAF) also suggest that they may drive T cell marginalization by forming different types of barrier to lymphocytes. While ECM degradation can improve T cell infiltration (7, 54), targeting ECM molecules is challenging in patients given the low specificity and the risk of on-target off-tumor toxicity. Here, our study paves the way to develop novel strategies to differently target the two distinct cellular sources of these ECM molecules.

In summary, our study has identified several CAF populations that show greater heterogeneity than previously established CAF classification, and provides novel therapeutic targets to pursue in order to augment response to cancer immunotherapies. We demonstrate that pairing molecular and spatial analysis is crucial to understanding the true organization of the human TME and to development of novel CAF targeting strategies for efficient anti-tumor combinations.

## SUPPLEMENTAL EXPERIMENTAL PROCEDURES

### Human subjects

In collaboration with the Biorepository and Department of Pathology tumor and adjacent non-involved lung samples were obtained from surgical specimens of patients undergoing resection at the Mount Sinai Medical Center (New York, NY). Informed consent was obtained in accordance with the following protocol reviewed and approved by the Institutional Review Board at the Icahn School of Medicine at Mount Sinai, IRB Human Subjects Electronic Research Applications 10-00472 and 10-00135.

### Tissue processing

The non-involved lung and tumor tissue were weighed and cut into sections of 0.1-0.2 grams then placed into 5 mL microtube (Argos Technologies). Sections were minced with scissors and enzymatically digested in CO2-independent media (Fisher Scientific, 18045088) with Collagenase IV 0.25mg/ml (Sigma-Aldrich, C5138-1G), Collagenase D 200 U/ml (Sigma-Aldrich, 11088882001), and DNAse 0.1 mg/ml (Sigma-Aldrich, DN25-1G) for 40 minutes at 37°C under 80 rpm agitation. Cell suspensions were passed through a syringe with an 18-gauge needle 8-10 times, filtered through a 70µm cell strainer, then lysed in red blood cell (RBC) buffer (Fisher Scientific, NC9067514). The cells were resuspended in buffer comprising of DPBS (Corning, D8537-6X500ML) with 5% BSA (Equitech-Bio, BAH62-0500) and 1mM EDTA (Sigma-Aldrich, 46-034-CI) then counted using hemocytometer and Trypan blue (Fisher Scientific, MT25900CI).

### Flow cytometry sorting

Cells were stained for EpCAM (Biolegend, clone 9C4), CD45 (Biolegend, clone HI30), CD29 (Biolegend, clone TS2/16), PDPN (Biolegend, clone NC-08), and LiveDead blue fluorescent dye (Thermo Scientific, L34963) for 30 minutes at 4°C. Among live cells, EpCAM and CD45 were used to remove epithelial and immune cells respectively, while CD29, present on all stromal cells, was used to enrich for cells with intact surface markers (see fig. S1B). The 1.5ml collection tubes (Fisher Scientific, 05-408-129) were coated with 10% BSA to improve cell survival post sorting.

### Single cell RNA sequencing

For each sample, up to an estimated 5,000 cells were loaded directly from the flow cytometry sort onto 10X Chromium chemistry kits. Kit versions for each sample are indicated in supplemental Table 1. Processing downstream of cell loading was performed by the Human Immune Monitoring Core at Icahn School of Medicine at Mount Sinai. Libraries were prepared according to manufacturer instructions and QC of the cDNA and final libraries was performed using CyberGreen qPCR library quantification assay. Sequencing was performed on Illumina sequencers to a depth of at least 80 million reads per library.

### Sequencing data analysis and unsupervised batch-aware clustering

Transcriptomic library reads were aligned to the GRCh38 reference genome and quantified using Cell Ranger (v3.1.0).

Stromal cells isolated from tumor and adjacent lung samples were analyzed using an unsupervised batch-aware clustering method we have recently described(30). First, stromal cells were filtered for cell barcodes recording > 800 UMI, with < 25% mitochondrial gene expression, and with less than defined thresholds of expression for genes associated with red blood cells, epithelial cells, macrophages, T cells and plasma cells (Table 12). 13 tumor and 11 adjacent samples were clustered jointly. This EM-like algorithm iteratively updates both cluster assignments and sample-wise noise estimates until it converges, using a multinomial mixture model capturing the transcriptional profiles of the different cell-states and sample specific fractions of background noise. We ran the algorithm described in Martin et al with minor modifications: Training and test set sizes per sample were 7500 and 2500 respectively. The best clustering initiation was selected from 1000 instead of 10000 k-means+ runs. For this clustering we included barcodes with more than 800 UMIs and used *K_reg_ds_* = 0.2; (P_1,_P_2_) = (0^th^,50^th^) percentiles; *K_reg_* = 5·10^−6^; k=28. Genes with high variability between patients across were not used in the clustering. Those genes consisted of mitochondrial, stress, metallothionein genes, immunoglobulin variable chain genes, HLA class I and II genes and 3 specific genes with variable/noisy expression*: MALAT1, JCHAIN* and *XIST* (Table 12). Ribosomal genes were excluded only from the k-means clustering (step 2.D as described in Martin et al.) (Table 12).

### Cell annotation

Using the gene module analysis described earlier, we identified highly variable genes and explored their expression across different clusters. Clusters were annotated by comparing gene expression patterns with profiles reported in prior literature.

For stromal cell clusters, EC expressed multiple identifying markers like *PECAM1*, *VWF*, *CLDN5*, and *EMCN* and lymphatics could be identified with *TFF3*, *LYVE1*, and *PROX1*. PvC were identified by combination of subset specific markers for and shared expression of contractile genes like *ACTA2*, *TAGLN*, *MYL9* and *TPM2*. PvC subset specific genes included *RGS5*, *COX4I2* and *HIGD1B* for pericytes and *DES* and *ACTG2* for SM. Identifying fibroblast markers included those listed in the main text, *PDGFRA*, *SPON1*, and *MMP2*, but also *DCN*, *FBLN1*, *LUM*, *COL1A2*, *RARRES2*, *CTGF*.

A cluster (#13) with contaminating epithelial cells was identified by the high expression of multiple keratin genes including *KRT17* and *KRT19*. Contaminating macrophages were identified in cluster 6 by expression of CD45 (*PTPRC*), *C1QB*, *C1QA*, *C1QC*, and *MARCO*. Cluster 18 and 16 were excluded due to high mitochondrial gene content and hemoglobin genes, respectively. The annotation process for fibroblast subsets is described in the text.

### Histological staining

Multiplexed IHC was performed according to the protocol developed by (28) with some modifications. Slides were baked at 37°C overnight, deparaffinized in xylene, then rehydrated. Antigen retrieval was done in citrate buffer (pH 6 or 9) (Dako, S2367 or 2369) at 95°C for 30 minutes, followed by incubation in 3% hydrogen peroxide for 15 minutes, then blocked using serum-free protein block solution (Dako, X0909) before adding primary antibody for 1 hour at room temperature or overnight at 4°C. The primary antibody was detected using a secondary antibody conjugated to horseradish peroxidase followed by chromogenic revelation using 3-amino-9-ethylcarbazole (AEC) (Vector laboratories, SK4200). Slides were counterstained with hematoxylin (Sigma-Aldrich, HHS32-1L) and mounted with a glycerol-based mounting medium (Dako, C0563). Then the same slides were bleached and re-stained as previously described. Antibodies sources can be found in Supplemental Table 4. Masson’s trichrome staining was performed by the Biorepository and Pathology core at the Icahn School of Medicine at Mount Sinai.

### Mass Cytometry by Time Of Flight (CyTOF)

Samples were processed to a single cell suspension according to the tissue processing protocol listed earlier. Cell viability staining was achieved with Rh103 staining for 20 minutes at 37°C followed by staining with the CyTOF antibodies listed in Supplemental Table 4. Acquisition of the samples was performed by the HIMC at mount Sinai. All analysis of the CyTOF samples, including the creation of viSNE plots, was done using the Cytobank platform (https://www.cytobank.org/).

### ECM and immunomodulatory gene expression profiles

Gene lists were sourced from (60) and (61) for ECM and immunomodulatory genes, respectively. Selected genes had to meet a mean expression threshold of 1 UMI per 2000 UMIs in 2% of cells in at least 1 cluster and meet a minimum 3-fold expression change between at least two clusters. Selected genes for display were further refined by qualitative analysis.

### Gene module analysis

Gene correlation modules were generated using a similar method as previously described in (30). Briefly, cells are downsampled to 2000 total UMIs and highly variable genes are isolated. A gene-gene correlation matrix for the isolated gene set is computed for each sample over the cell population(s) of interest and correlation matrices are averaged following a Fisher Z-transformation. Applying the inverse transformation then results in the best-estimate correlation coefficients of gene-gene interactions across the dataset. Genes are clustered into modules using complete linkage hierarchical clustering over correlation distance. Ribosomal, mitochondrial, HLA and immunoglobulin genes were removed from the analysis prior to creation of gene modules as these genes were not of interest in this study, reflected patient genomic variability or were heavily influenced by contaminating plasma cells.

### Acquisition of TCGA dataset and histological subtypes

The TCGA lung adenocarcinoma (LUAD) RNAseq data was downloaded using the *GDCquery* and *GDCdownload* functions from the *TCGAbiolinks* R package. *GDCquery* options included *project=”TCGA-LUAD”, data.category=”Transcriptome Profiling”, data.type=”Gene Expression Quantification”, workflow.type=”HTSeq – FPKM”, experimental.strategy=”RNA-Seq”,* and *legacy=F*. Whole exome sequencing data was downloaded using the *GDCquery_Maf* function with arguments *tumor=”TCGA-LUAD”* and *pipelines=”mutect2”*. Clinical data was downloaded using the *GDCquery_clinic* function with arguments *project=”TCGA-LUAD”* and *type=”clinical”*.

The dominant histological subtype for each TCGA tumor was sourced from (42).

### CAF gene signatures and LCAM in bulk analysis

We performed gene module analysis to identify groups of co-expressed genes in the scRNAseq dataset as described above. To define cell type specific gene signatures, we first excluded genes expressed in non-fibroblast lineages, such as EC, PvC, epithelial, and immune cells. We then compared the gene module expression between fibroblast subsets. Due to their similarity, FAP^+^ CAF and FAP^+^αSMA^+^ CAF were treated as one group and due to ADH1B^+^ CAF similarity to adjacent fibroblast clusters they were not contrasted with alv. fib. and *PI16*^+^ fib. Signatures were further refined by manually checking that expression was consistently enriched in the cell type of interest across at least 3 patients.

Bulk RNA gene expression values were log-transformed and z-scored. Cell type signatures, scored per sample, were calculated by taking the average of all genes within the signature. The derivation of LCAM^high^ or LCAM^low^ is described in [Leader et al. 2021]. The cell types associated with each state were averaged to create an LCAM^high^ or LCAM^low^ score. The difference between LCAM^high^ and LCAM^low^ was the final LCAM score. All signatures are calculated using only tumor samples, with sample ID ending in ‘-01A’, and signatures were z-scored before graphing or other analysis. TCGA patients with their corresponding signatures scores can be found in Table 6.

### Histology analysis and overlays

All histology analysis was performed using the open-source image analysis QuPath software (QuPath-0.2.3, https://qupath.github.io/)(62) and ImageJ/Fiji(63, 64).

To create overlayed images, scans exported from QuPath as OME.TIFF then imported into ImageJ using the BioFormats plugin(65). Alignment was done using the “Linear Stack Alignment with SIFT” plugin(66). The AEC and hematoxylin stains were extracted from individual scans using “colour deconvolution” and colored as desired.

#### Quantifying T cell infiltration and inter-patient CAF heterogeneity

Cropped scans were imported into a newly created project in QuPath and were aligned using the “interactive image alignment” plugin. Alignment information was saved using QuPath_script_1. Tumor and stroma regions were defined by applying the “create cytokine annotation” function on the keratin scans. The stroma and tumor annotations were transferred onto the aligned CD3, CD8, αSMA, ADH1B, and FAP scans with QuPath_script_2. On the CD3 and CD8 scans, positive cell detection was used to count the CD3^+^ and CD8^+^ cells. Distance to the tumor and stroma annotations was calculated using the “distance to annotation 2D” option and the measurements were exported as raw data to be analyzed in R. For figure 4E the stroma and tumor annotations were tiled using QuPath function “Create Tiles” and αSMA, ADH1B, and FAP scans the positively stained area were calculated using the QuPath training classifier.

#### Quantifying T cell infiltration and intra-patient CAF heterogeneity

For the intra-patient analysis, we created a separate QuPath project with all the desired scans to analyze. The images were cropped and exported and then overlays were generated using an ImageJ script with the following steps:

a. Deconvolution (hematoxylin, AEC, residual)
b. Alignment on hematoxylin images
c. Creation of transformation matrix then application on AEC images
d. Threshold to remove background and then ”Stack of Images”

The composite image was transferred back to QuPath for further analysis. Adjacent tissue regions on slides were excluded from the analysis and in the region to analyze we used the “train pixel classifier” to annotate Tumor nests versus Stroma; while αSMA, FAP and ADH1B areas were annotated using “QuPath train classifier”. Dedicated script automated the tiling and quantification and resulted in 3 data files; cell information including staining intensity and distances to annotations, tile annotation parameters, and annotation measurements such as ADH1B stained area within an annotation. The resulting measurements were exported and analyzed in R.

#### Quantifying MYH11^+^αSMA^+^ CAF boundary enrichment

MYH11^+^αSMA^+^ CAF score (Figures 5C, 5D) approximates the enrichment of these cells at the tumor nest boundary (<10μm) relative to their distal background density (20-30μm). Iterating over tiles, we counted the number of MYH11^+^αSMA^+^ double positive cells in each distance bin of 1μm. The proximal value was defined as the quantile, 0.75, of the number of cells in the bins within distance <10μm. The distal value was similarly defined as the quantile, 0.75, of the number of cells in the bins with distances between 20-30μm. The MYH11^+^αSMA^+^ CAF score was defined as log2(proximal/distal). The 0.75 percentile was selected to maximize the sensitivity of detecting robust high-density regions.

## DATA AVAILABILITY

Sequencing data will be available through GEO and accession number will be provided by publication date. Code is available on our Github website: https://github.com/effiken/Grout_et_al.

## ACKNOWLEDGMENTS

The authors thank the important contributions of patients who participated in this study. This project was supported by Genentech, Inc. and carried out in collaboration with the Fondation ARC pour la recherche sur le cancer. The computational work was supported by the Scientific Computing at the Icahn School of Medicine at Mount Sinai and the Office of Research Infrastructure of the National Institutes of Health under award number S10OD026880. The content is solely the responsibility of the authors and does not necessarily represent the official views of the National Institutes of Health. This manuscript was edited at Life Science Editors and the cartoon illustrations were created using BioRender.com. We thank the Mount Sinai Flow Cytometry core, Human Immune Monitoring Center and the Biorepository and Pathology core for their support. We thank Eliane Piaggio, Ana-Maria Lennon-Duménil, and Olivier Lantz for carefully reading and commenting the manuscript.

## AUTHOR CONTRIBUTIONS

HS, and EK conceived the project. JAG, FS, NT, and HS designed the experiments. JAG, EK, and HS wrote the manuscript. JAG, PS, AML, EH, RP, SMü and EK performed computational analysis. AW, MBB, and RF facilitated access to human samples. JAG, SMa, IP, EC, NT, MM, SK, RV, LK, MP, MCA, AT and LW performed experiments and other analysis. AL conducted patient consents and facilitated regulatory items. MM, and SJT provided key guidance. AML, AHR, SG, JA, JCM, MBB, SMü, and TUM provided further intellectual input.

## SUPPLEMENTAL TABLES

S1 – Patient clinical summary data

S2 – Patient cell counts per cluster

S3 – Adjacent and tumor sample percent composition of fibroblast clusters

S4 – Antibodies used in IHC, flow, and CyTOF

S5 – Positive staining area of ADH1B and FAP for IHC slides

S6 – TCGA patient expression of signatures

S7 – CD8 and CD3 quantification across entire FFPE section

S8 – Slide tiling data for FAP^+^αSMA^+^ CAF – sample 24

S9 – Slide tiling data for FAP^+^αSMA^+^ CAF – sample 54

S10 – Slide tiling data for MYH11^+^αSMA^+^ – CAF sample 13

S11 – Slide tiling data for MYH11^+^αSMA^+^ – CAF sample 8

S12 – Gene lists used in scRNAseq clustering, *in silico* filtering and in figures

S13 – CD8^+^ T cell tumor density across NSCLC samples

