## Supplemental figures for "Spatial positioning and matrix programs of cancer-associated fibroblasts promote T cell exclusion in human lung tumors"

### Figure S1, associated with figure 1

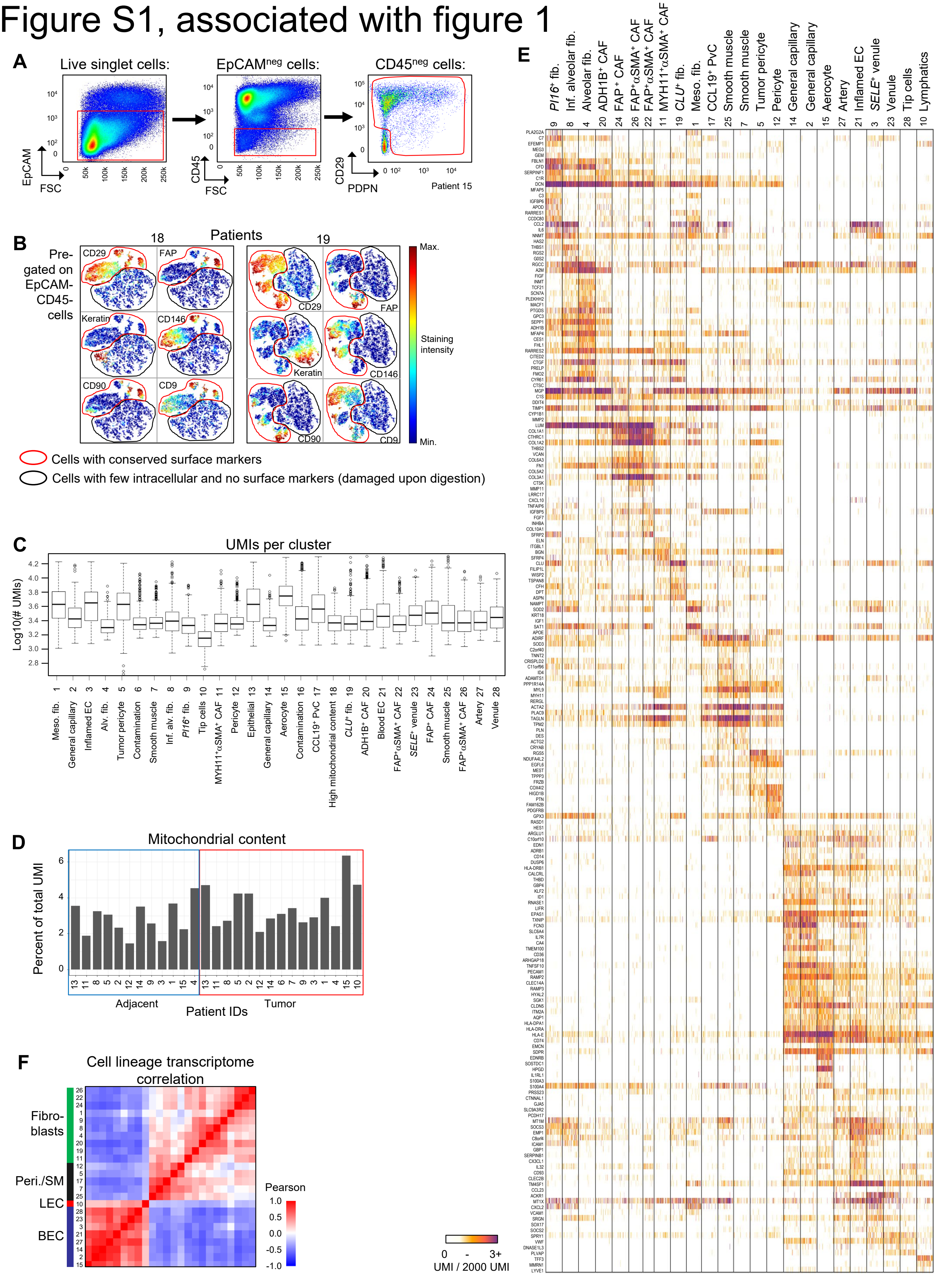

**Supplemental fig. 1 (related to fig. 1) | A**, Flow cytometry sorting strategy to enrich for stromal cells. EpCAM and CD45 gating removed epithelial and immune cell populations, respectively. Integrin β1 (CD29) and podoplanin (PDPN) positive gating was used to remove any remaining damaged cell that lost surface markers upon tissue digestion (as seen in Fig. S1B). **B**, CyTOF viSNE representation of pre-gated cisplatin$^{neg}$ (for dead cell exclusion) EpCAM$^{neg}$ CD45$^{neg}$ cells obtained from tumor lesions of two NSCLC patients (#18 and 19). The two viSNE plots show a substantial cell cluster (circled in black) lacking the 15 surface markers in the CyTOF antibody panel (here, 5 surface markers – CD29, FAP, CD146, CD90, CD9 – are shown as representative examples), while preserving levels of intracellular markers (such as keratin and vimentin). These results indicate that these cells likely contain live but damaged cells that lost surface protein expression upon tissue digestion (potentially tumor, immune, or stromal damaged cells). They could also contain red blood cells. Notably, all remaining cells expressed CD29, which was therefore selected as a positive marker including all undamaged stromal cells. **C**, Boxplot of UMI counts per scRNAseq cluster. **D**, Mitochondrial gene content for all sequenced cells in the tumor and its adjacent stroma for each patient. **E**, Highly variable genes across all stromal clusters. Genes were selected if they were highly variable (Log2(var/mean) = 1.25) and met a minimum expression cutoff of 8 UMI across all cells included. **F**, Heatmap showing the correlation (Pearson) between whole genome expression profiles of major stromal populations. Fibroblast, perivascular, and endothelial lineages show high concordance as expected. Gene expression values per cluster are determined by taking the mean expression of each gene across all cells then taking the log2 value.

### Figure S2, associated with figure 1

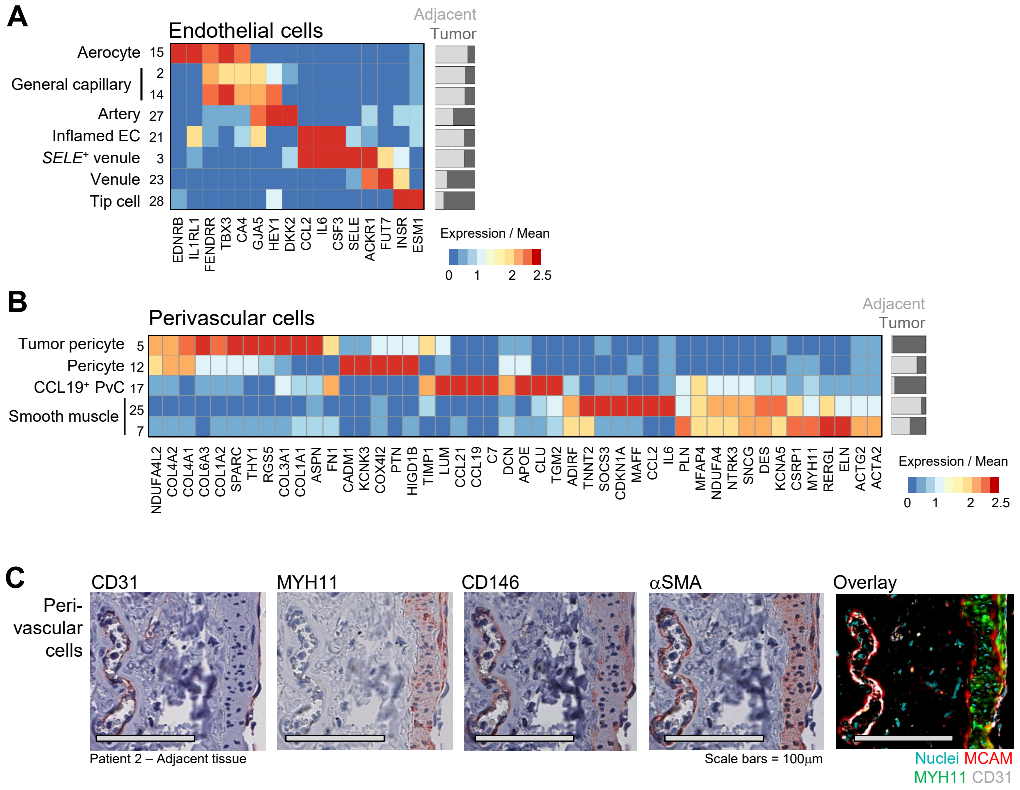

**Supplemental fig. 2 (related to fig. 1) | A-B**, (**A**) Gene lists highlighting identifying genes and heterogeneity between EC clusters and (**B**) PvC clusters. **C**, IHC staining of blood vessel showing pericytes (CD146 and αSMA), smooth muscle cells (CD146, αSMA and MYH11) and ECs (CD146 and CD31). All scale bars are 100μm.

### Figure S3, associated with figure 1

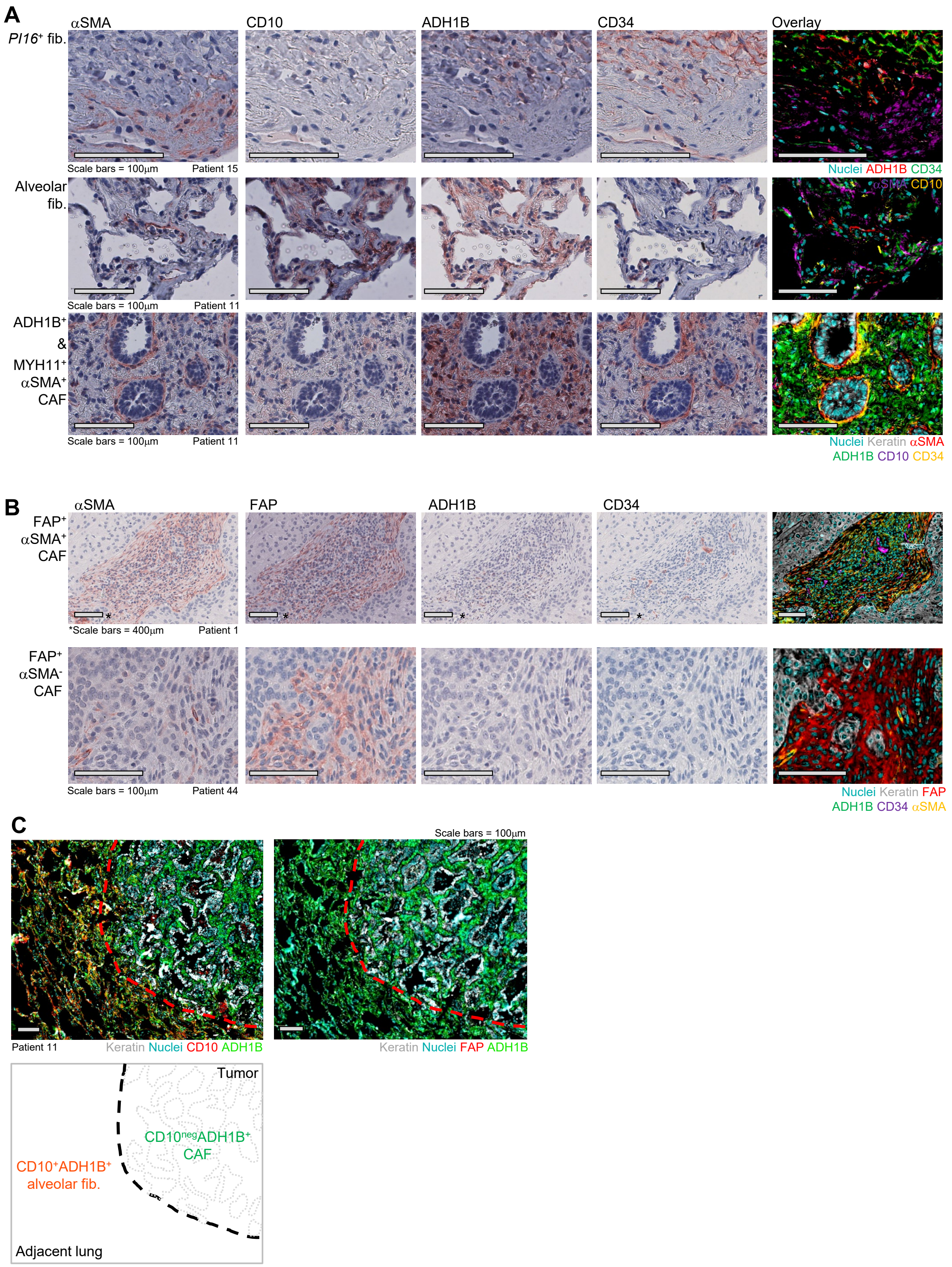

**Supplemental fig. 3 (related to fig. 1) | A-B**, Component IHC stains of MICSSS panels. (**A)**, IHC stains highlighting *PI16*$^{+}$ fib., alv. fib., ADH1B$^{+}$ CAF and MYH11$^{+}\alpha$SMA$^{+}$ CAF. All scale bars are 100μm. (**B)**, IHC staining of FAP$^{+}$ CAF highlighting heterogeneity within this fibroblast subset demonstrated by αSMA intensity. FAP$^{+}$ CAF (top panels) scale bars are 100μm and FAP$^{+}\alpha$SMA$^{+}$ CAF scale bars are 400μm. **C**, Multiplexed IHC highlighting lack of FAP and CD10 staining in ADH1B$^{+}$ CAF cells. All scale bars are 100μm.

### Figure S4, associated with figure 3

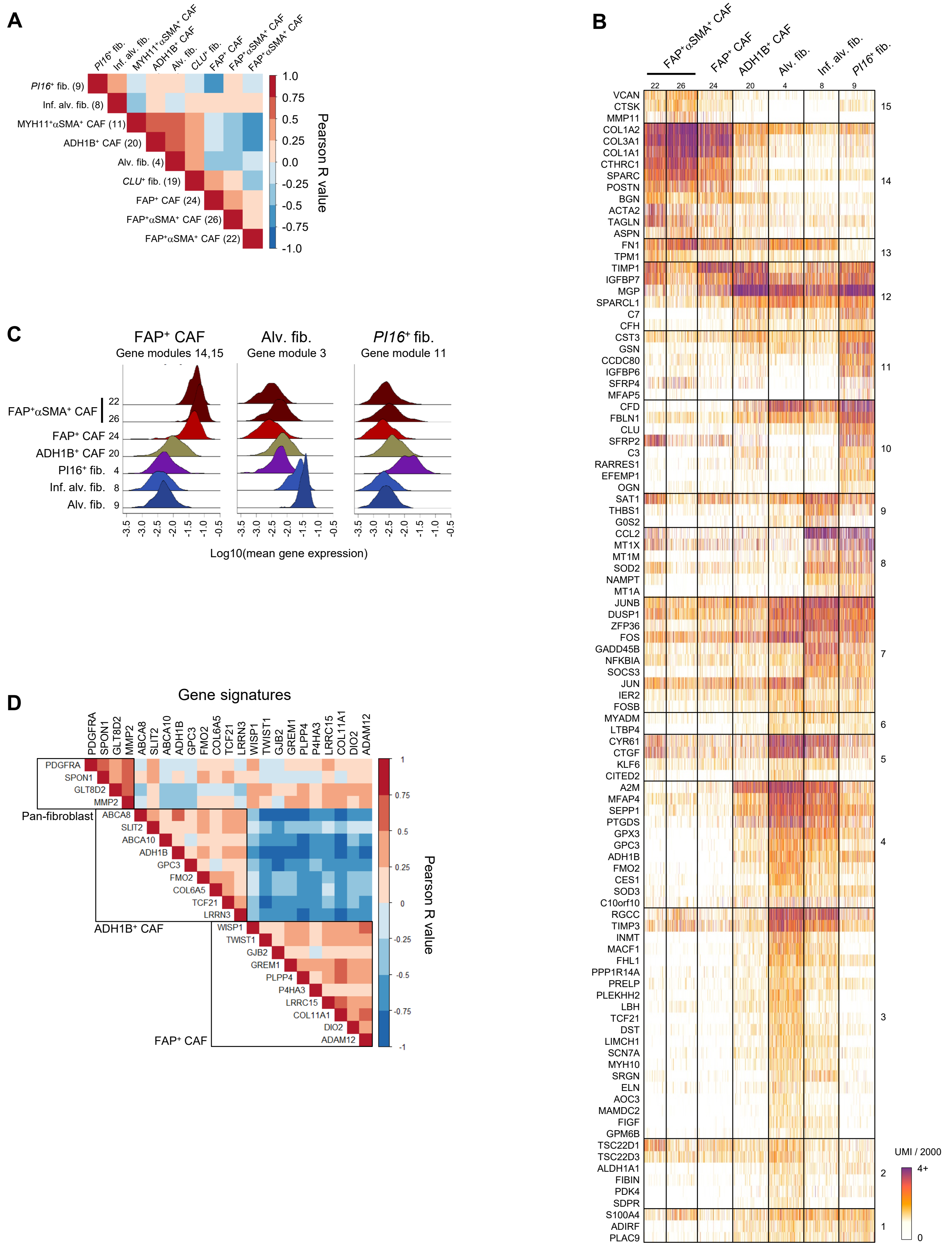

**Supplemental fig. 4 (related to fig. 3) | A**, Correlation (Pearson) of fibroblast subset abundance in scRNAseq tumor samples. **B**, Co-expressed gene 'modules' identified within the fibroblast clusters. Each module contains a group of genes with highly correlated expression regardless of enrichment in any particular cluster. These modules frequently represent identifying genes for specific clusters, such as modules 14 and 15 for $FAP^+$ CAF, but may also represent groups of genes co-regulated across multiple clusters, like module 7. Genes were selected if they were highly variable (Log2(var/mean) = 0.9) and met a minimum expression cutoff of 10 UMI across all cells included. **C**, Mean log10 gene expression of an $PI16^+$, alv. fib. and $FAP^+$ CAF-associated gene modules displayed by density histogram. Higher peaks indicate greater cell numbers at the expression level indicated on the X axis. Single-cell gene expression of modules is shown in Figure S4B. **D**, Correlation (Pearson) of $ADH1B^+$ CAF and $FAP^+$ CAF gene signature components in TCGA.

### Figure S5, associated with figure 5

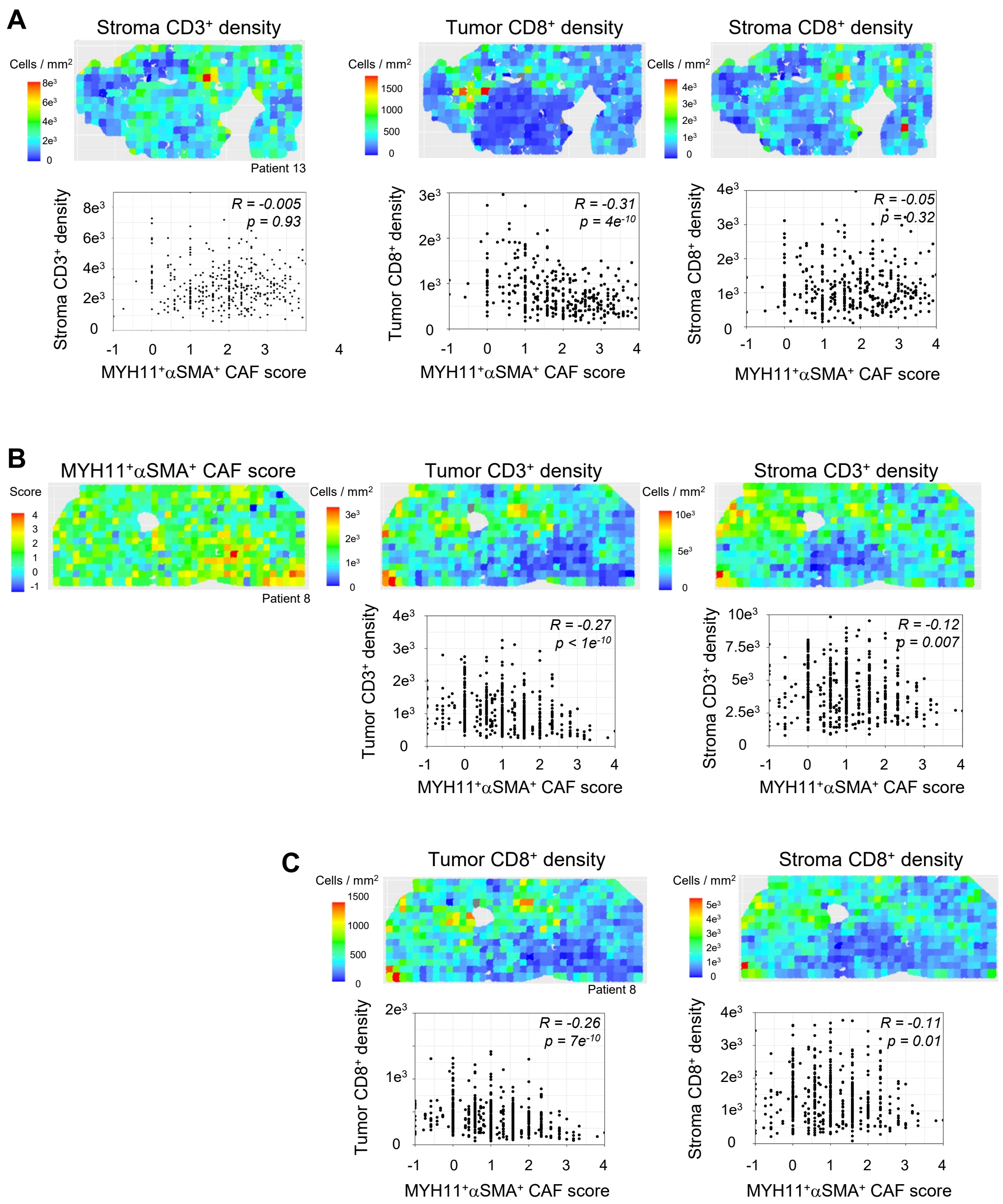

**Supplemental fig. 5 (related to fig. 5) | A**, Heatmaps of $CD3^+$ cell density in the stroma and $CD8^+$ cell density in the tumor and stroma of the patient 13. Respective dot plots and correlation (Pearson) with $MYH11^+\alpha SMA^+$ CAF score shown below each heatmap. **B**, $MYH11^+\alpha SMA^+$ CAF score, $CD3^+$ cell density heatmaps in the tumor and stroma of the patient 8. Dot plots of $CD3^+$ cell density correlation (Pearson) with $MYH11^+\alpha SMA^+$ CAF score shown below each heatmap. **C**, $CD8^+$ cell density in the tumor and stroma of the patient 8. Dot plots of $CD8^+$ cell density correlation (Pearson) with $MYH11^+\alpha SMA^+$ CAF score shown below each heatmap.

### Figure S6, associated with figure 6

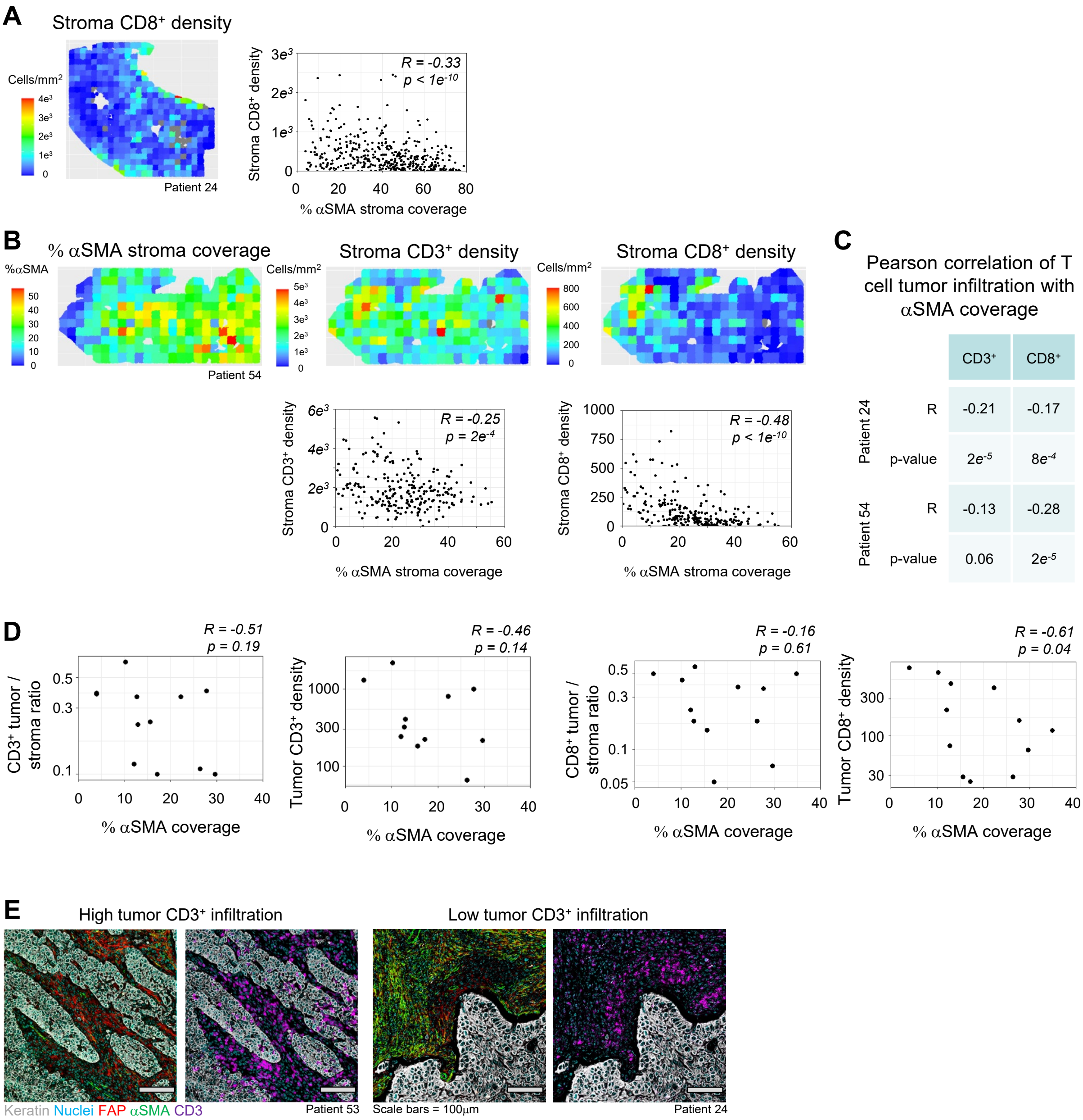

**Supplemental fig. 6 (related to fig. 6) | A**, Stroma $CD8^+$ cell density heatmap of patient 24 (left panel) and corresponding correlation (Pearson) dot plot with αSMA stroma coverage (right panel). **B**, αSMA coverage, stroma $CD3^+$ and $CD8^+$ cell density heatmap of the patient 54 (top panels) and corresponding correlation (Pearson) dot plots of $CD3^+$ and $CD8^+$ cell density with αSMA stroma coverage (bottom panels). **C**, Correlation (Pearson) of $CD3^+$ and $CD8^+$ cell tumor infiltration with stroma αSMA coverage in patients 24 and 54. **D**, Comparison of αSMA stroma coverage with $CD3^+$ and $CD8^+$ cell tumor infiltration for each patient. αSMA coverage and tumor infiltration is quantified across the entire tumor lesion. **E**, Example tumor lesions displaying high or low $CD3^+$ cell infiltration.
